# A bacterial nucleolus for cold adaptation

**DOI:** 10.64898/2026.09.17.752289

**Authors:** Michelle J. Gut, K. Dörner, Sebastian Abegg, Nusrat S. Qureshi, Kausthubh Ramachandran, Alexia Ferrand, Alexander Schmidt, Ruta Prakapaite, Tom A. Williams, Olivier Duss, Maria Hondele

## Abstract

Compartmentalisation enables cells to spatially organise essential biochemical processes. This is exemplified by the eukaryotic nucleolus, which concentrates the machinery for ribosome biogenesis^1,2^. Bacteria are generally thought to lack an equivalent compartment. Here we discover a nucleolus-like condensate in *Escherichia coli* that emerges at rDNA loci during cold adaptation to spatially organise and promote ribosome biogenesis. Low temperatures, which stabilise RNA secondary structures^3–8^, induce accumulation of the RNA-remodelling enzyme CsdA^9,10^. This raises the cellular CsdA concentration above its condensation threshold, driving condensate formation at rDNA loci when rRNA transcription is active. CsdA compartments enrich the rRNA transcription machinery and ribosome-biogenesis factors, spatially linking rRNA synthesis to downstream processing and assembly. Disrupting CsdA condensation impairs ribosome maturation and reduces bacterial adaptation to cold temperature, demonstrating functional importance of condensate formation. Comparative genomics suggests that analogous condensate-forming RNA remodelling enzymes have evolved repeatedly across diverse bacterial lineages. Our findings establish biomolecular condensates as a unifying organisational principle of ribosome biogenesis across the tree of life and suggest that adaptive compartmentalisation can emerge when environmental conditions challenge the efficiency of essential biochemical processes.

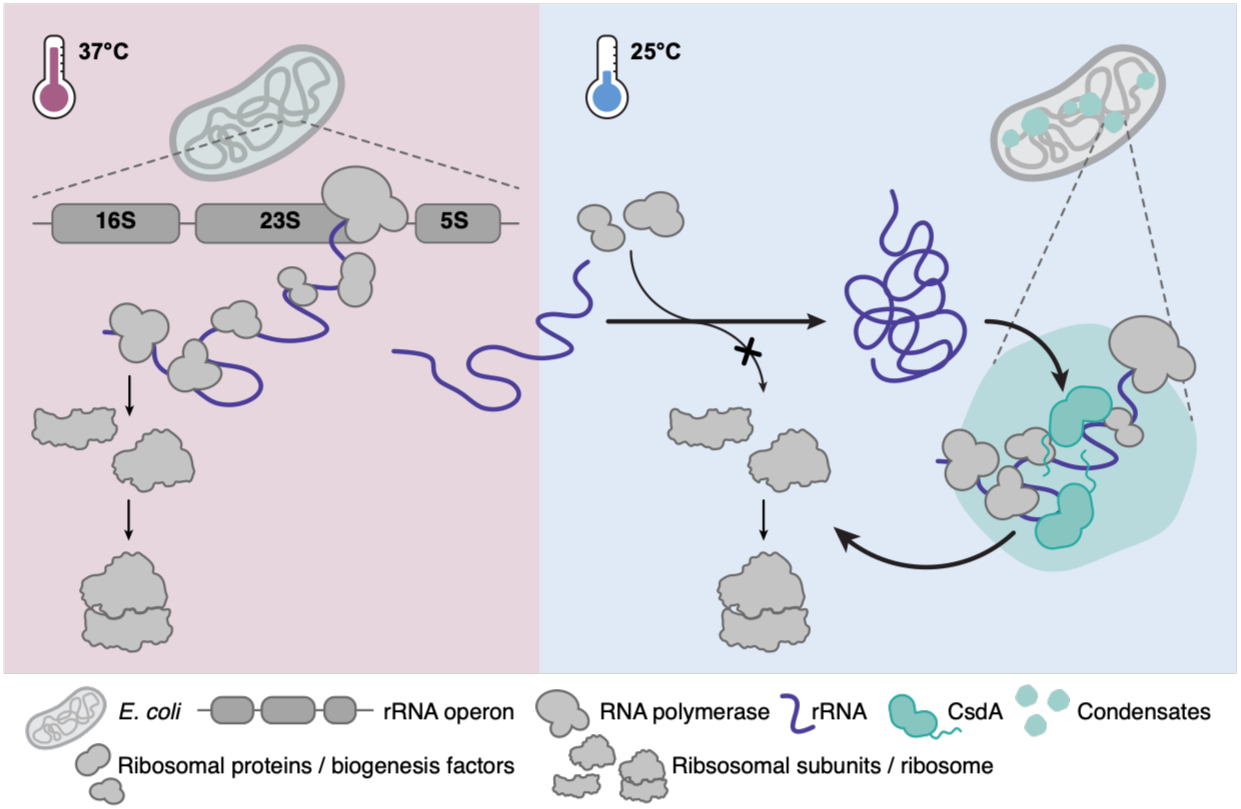

## Main Text

Ribosome production requires the coordinated transcription, processing and folding of ribosomal RNA (rRNA), and its assembly with ribosomal proteins. In eukaryotes, these reactions are spatially organised within the nucleolus, a multilayered biomolecular condensate that couples rRNA synthesis to successive stages of processing and ribosome assembly^1,2^. Fast-growing bacteria assemble transcription factories in which RNA polymerase and antitermination factors cluster at highly transcribed rRNA operons^11–14^, yet these assemblies are understood primarily as sites of rRNA synthesis, whereas downstream ribosome assembly is generally thought to proceed without a dedicated compartment.

Environmental conditions can, however, challenge ribosome biogenesis. Low temperatures stabilise RNA secondary structures and trap nascent rRNA in non-productive conformations^3–8^. Bacteria counteract RNA misfolding by upregulating several RNA-remodelling enzymes, including the highly conserved family of DEAD-box ATPases (DDXs)^9,10,15^. *Escherichia coli (E. coli)* encodes five DDXs, of which CsdA/DeaD is the major cold-induced enzyme and promotes rRNA folding and ribosomal-protein incorporation at low temperature^9,16,17^.

Beyond their enzymatic functions, eukaryotic DDXs form and organise RNA-rich condensates, including the nucleolus and stress granules^1,18–23^. Several bacterial DDXs likewise condense in vitro or when overexpressed in cells^18,24,25^. Whether bacteria exploit DDX condensation under physiological conditions to spatially coordinate RNA remodelling and in particular ribosome biogenesis remains unknown.

### CsdA forms regulated foci during cold adaptation

To understand how bacteria adapt rRNA processing to low temperatures and whether it involves the assembly of specialised RNA-remodelling condensates, we examined the localisation of the five *E. coli* DDXs. Each DDX was endogenously tagged with mEGFP and visualised by three-dimensional structured illumination microscopy (3D-SIM) in cells grown at optimal growth temperature (37 °C) or low temperature (25 °C) (Fig. 1a-c and Extended Data Fig. 1a-c). RhlE and DbpA remained diffusely distributed under both conditions, whereas SrmB and RhlB formed temperature-independent foci both at 37 °C and 25 °C. SrmB foci localised to the nucleoid, whereas RhlB formed peripheral foci likely corresponding to membrane-associated RNA decay compartments (BR bodies) (Extended Data Fig. 1b)^26^. Interestingly, only CsdA underwent a pronounced temperature-induced redistribution into prominent nucleoid-associated foci at low temperature (Fig. 1a-c, Extended Data Fig. 1a-c).

**Figure 1.**
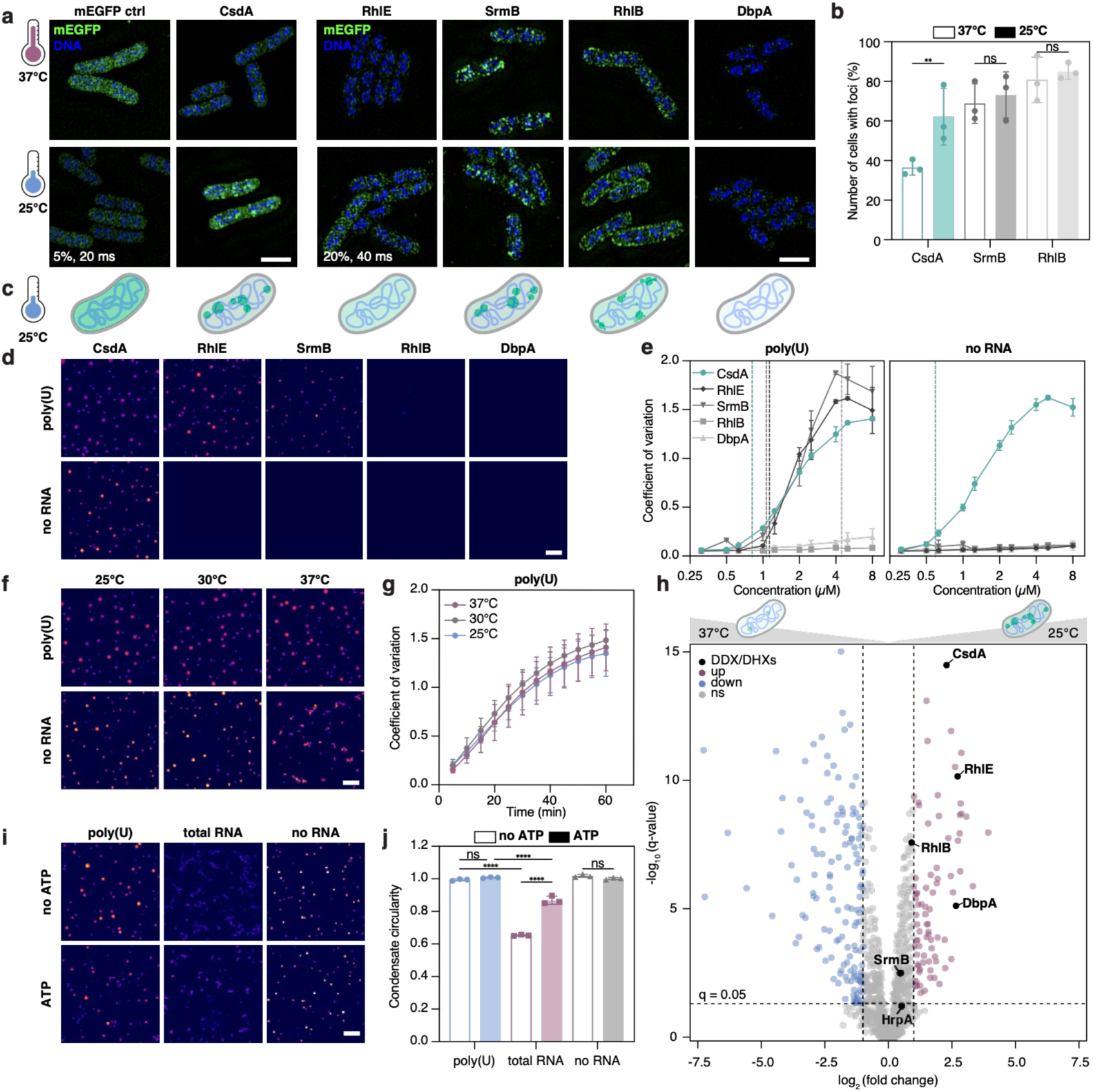
Cold-induced CsdA foci emerge when CsdA crosses its condensation threshold. **a,** 3D-SIM of the five endogenously mEGFP-tagged *E. coli* DDXs and soluble mEGFP at 37°C and 25°C; DNA was stained with Hoechst. Due to large differences in signal intensity, CsdA and mEGFP control were imaged at lower exposure (5%, 20 ms), and other DDXs at higher exposure (20%, 40 ms), all images shown with the intensity range (N=3), scale bar 2 µm. **b,** Percentage of cells containing foci. **c,** Schematic of DDX localisation at 25°C. **d,** Representative condensation of 2 μM recombinant DDXs ± poly(U) at 25°C (N=3), for all in vitro condensation experiments scale bars 20 μm. **e,** Concentration-dependent quantification by fluorescence coefficient of variation; vertical dashed-lines indicate *c*_sat_. **f,** Representative condensation of 2 µM CsdA at 25°C, 30°C and 37°C ± poly(U) (N=3). **g,** Temperature-dependent condensation kinetics of 2 µM CsdA at 25°C, 30°C and 37°C with poly(U) (N=3). **h,** Relative quantitative proteomics of DDX abundance at 25°C versus 37°C; dashed lines indicate q=0.05 and |log_2_(FC)|=1 thresholds; blue, decreased; red, increased proteins at 25°C (N=3). **i,j,** CsdA condensation (2 µM) with poly(U), total *E. coli* RNA or no RNA ± ATP/ATP-regeneration system and condensate circularity quantified by form factor (N=3). **b,e,g,j,** Mean ± SD, individual replicates shown as dots. Significance assessed by ordinary one-way ANOVA.

### Cold-induced CsdA accumulation drives formation of CsdA condensates

The cold-induced appearance of CsdA foci and their sensitivity to 1,6-hexanediol (Extended Data Fig. 1d,e) indicated that CsdA foci are biomolecular condensates. The mechanism behind CsdA condensation during cold adaptation, however, remained unclear. We therefore compared the intrinsic condensation behaviour of purified recombinant CsdA and the other *E. coli* DDXs in vitro at 25 °C (Extended Data Fig. 2a,b). In the presence of RNA, CsdA, RhlE and SrmB formed condensates, whereas RhlB and DbpA remained dispersed (Fig. 1d and Extended Data Fig. 2c). Among these proteins, CsdA exhibited the lowest saturation concentration (*c*_sat_ ≈ 0.8 µM) and readily condensed even in the absence of RNA (*c*_sat_ ≈ 0.6 µM) (Fig. 1e, and Table 1). Thus, the only *E. coli* DDX forming cold-induced foci in cells also possesses the highest intrinsic in vitro condensation propensity.

**Table 1.** Cellular abundance and condensation thresholds of bacterial DEAD-box ATPases. Protein copy numbers and estimated intracellular concentrations *c*_cell_ at 37°C and 25 °C, proteomics-derived log_2_(fold change), and fold changes upon temperature shift, and experimentally determined in vitro *c*_sat_ in the presence or absence of RNA are shown for the five *E. coli* DEAD-box ATPases at 25 °C.

| Protein | 37°C (in cells) | | $\log_2(\text{fold change})$ | 25°C (in cells) | | 25°C (in vitro) | | | Expected cellular foci |
| --- | --- | --- | --- | --- | --- | --- | --- | --- | --- |
| | copies/cell | $c_{\text{cells}}$<br>($\mu\text{M}$) | | fold change | copies / cell | $c_{\text{cells}}$<br>($\mu\text{M}$) | $c_{\text{sat}}$ (poly(U))<br>($\mu\text{M}$) | $c_{\text{sat}}$ (no RNA)<br>($\mu\text{M}$) | |
| CsdA/DeaD | 1899 | 0.79 | 2.29 | 4.89 | 9285 | 3.85 | 0.81 (+/- 0.01) | 0.59 (+/- 0.03) | yes |
| RhlE | 54 | 0.02 | 2.73 | 6.63 | 359 | 0.15 | 1.13 (+/- 0.08) | no LLPS | no |
| SrmB | 719 | 0.30 | 0.47 | 1.39 | 996 | 0.41 | 1.07 (+/- 0.25) | no LLPS | no |
| RhlB | 938 | 0.39 | 0.91 | 1.88 | 1763 | 0.73 | > 10 $\mu\text{M}$ | no LLPS | no |
| DbpA | 28 | 0.01 | 2.66 | 6.32 | 176 | 0.07 | no LLPS | no LLPS | no |

Given CsdA’s high intrinsic condensation propensity, the pronounced increase in CsdA condensates during cold adaptation suggested that its condensate formation is regulated in vivo. Two non-mutually exclusive mechanisms appeared plausible. Low temperature could directly enhance CsdA phase separation, as described for other proteins^27,28^, but CsdA in vitro condensation kinetics and threshold were highly similar at 37 °C, 30 °C, and 25 °C (Fig. 1f,g, and Extended Data Fig. 2d,e). Alternatively, cold adaptation could upregulate cellular CsdA levels and thereby drive CsdA across its condensation threshold. Proteomics experiments indeed revealed a significant increase in CsdA abundance at 25 °C (fold change = 4.9) (Fig. 1h and Extended Data Fig. 1f,g). Based on previously reported protein copy numbers per cell at 37 °C and an *E. coli* cell volume of ∼4 fl^29,30^, we estimated the intracellular CsdA concentration to increase from 0.8 µM at 37 °C to 3.9 µM at 25 °C, thereby raising CsdA above its *c*_sat_ (Table 1). By contrast, the other *E. coli* DDXs remained well below their respective saturation concentrations.

Thus, CsdA condensation is primarily regulated by tuning protein abundance rather than its phase behaviour. CsdA concentration is maintained close to *c*_sat_ during optimal growth, allowing a modest increase in abundance during cold adaptation to raise its cellular concentration above the saturation threshold.

### CsdA condensates locally concentrate RNA remodelling activity

Low temperatures stabilise RNA secondary structures and can trap RNA in non-native conformations, creating folding barriers that impair processes such as ribosome biogenesis^3,4,7,31^. If CsdA condensation represents an adaptive response to cold, it could create a local environment that facilitates RNA remodelling and helps overcome these folding barriers.

We therefore asked whether structured RNA drives changes in CsdA condensate morphology and whether CsdA’s ATP-dependent RNA remodelling activity can reverse this transition by resolving the underlying RNA-RNA interactions. To address this, we examined how CsdA catalytic activity reshapes condensates of different RNA structural complexity at 25 °C. Condensates assembled without RNA or with poly(U), a largely unstructured RNA, are round and dynamic. *E. coli* total RNA (Extended Data Fig. 2f), which contains persistent RNA-RNA interactions, shifted CsdA condensates towards more irregular, aggregate-like structures, indicated by the significantly lower circularity (Fig. 1i,j and Extended Data Fig. 2g,h). Addition of ATP, which activates CsdA-dependent RNA remodelling^32,33^, partially restored the roundish condensate morphology and rescued circularity, indicating that ATP-dependent RNA remodelling by CsdA counteracts the transition to aggregates.

This suggests that CsdA condensates locally concentrate RNA-remodelling activity, thereby counteracting persistent RNA–RNA interactions that become increasingly problematic at low temperature.

### CsdA condensates assemble at *rrn* operons and recruit ribosome biogenesis factors

If CsdA condensates create a locally concentrated environment for RNA remodelling, they might also recruit factors involved in rRNA synthesis, maturation, and ribosome assembly. Affinity purification–mass spectrometry (AP–MS) of CsdA-mEGFP from cells grown at 25 °C recovered numerous ribosome biogenesis factors and ribosomal proteins as well as the RNA polymerase subunits RpoA, RpoB, and RpoC (Fig. 2a). By contrast, RNA degradation factors were only weakly enriched, arguing against a primary role for these compartments in RNA decay^34^. Instead, the interaction landscape hints at a spatial coupling of rRNA synthesis, remodelling, maturation and ribosome assembly.

**Fig. 2.**
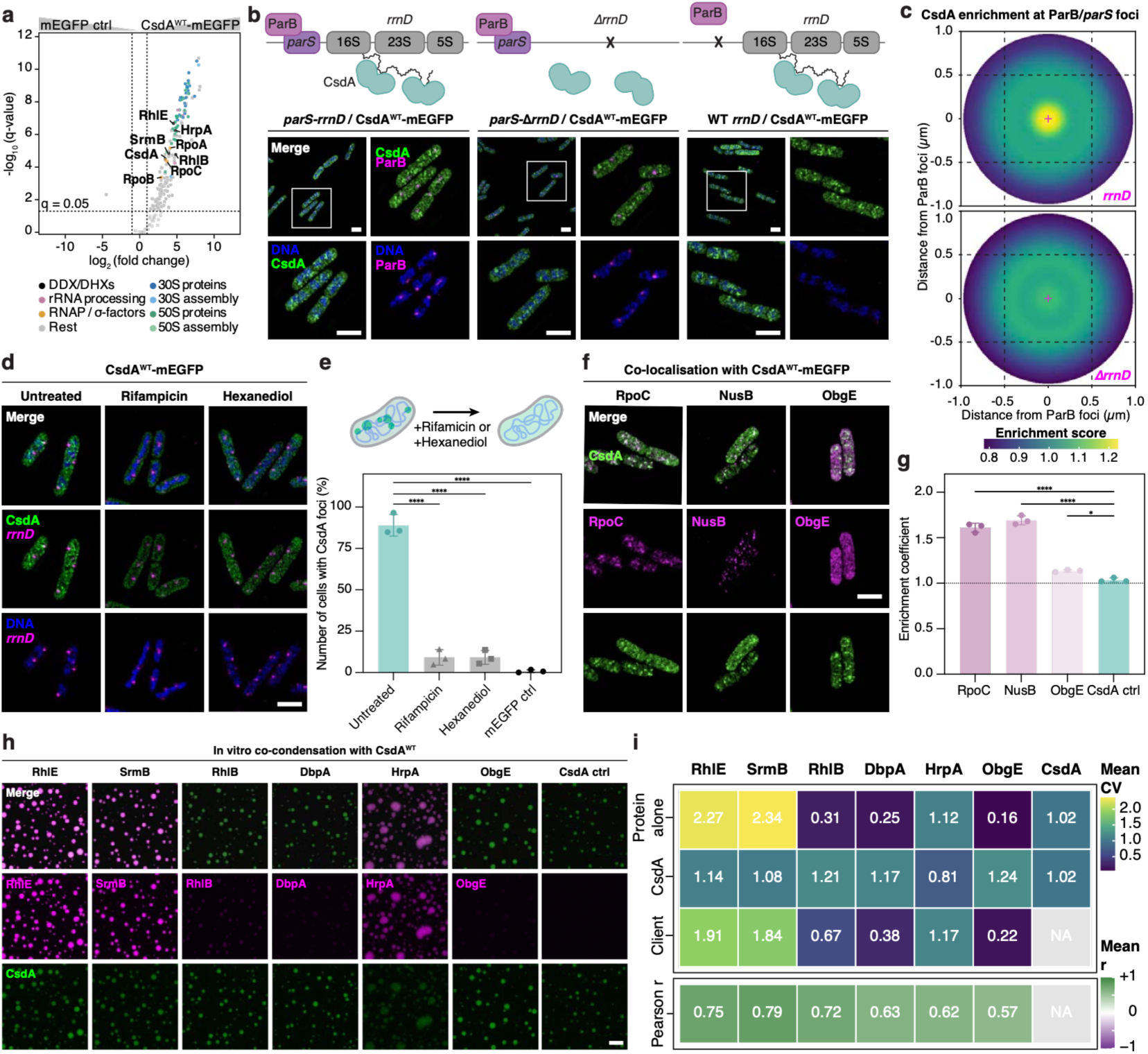
CsdA condensates associate with *rrn* loci and ribosome biogenesis factors. **a,** AP-MS with schematics of CsdA^WT^ interaction partners following cold adaptation relative to mEGFP at 25°C, highlighting DDXs, ribosomal proteins, ribosome biogenesis factors, RNA-processing factors, RNAP subunits and sigma factors (N=3). **b,** 3D-SIM of CsdA-mEGFP at 25°C in strains with mCherry-ParB/*parS*-marked *rrnD* locus; *ΔrrnD* and no *parS* control strains. DNA stained with Hoechst (N=3). For all 3D-SIM images: scale bar 2 µm. **c,** Radial analysis of CsdA foci mean distance from ParB/*parS* loci. **d,e,** CsdA localisation and percentage of cells containing foci following rifampicin or 1,6-hexanediol treatment at 25°C (N=3). **f,g,** Colocalisation and enrichment of plasmid-expressed RpoC, NusB and ObgE at CsdA condensates at 25°C (N=3). **h,** In vitro partitioning of recombinant ribosome-associated factors into CsdA condensates formed with poly(U) at 25°C (N=3). Scale bar 20 µm. **i,** Condensation quantified by mean coefficient of variation (CV) for individual proteins and co-condensation. and mean Pearson correlation of CsdA and client fluorescence within segmented CsdA condensates. **e,g,i,** Mean ± SD shown as bar graph, individual replicates shown as dots. Significance assessed by ordinary one-way ANOVA.

The enrichment of RNA polymerase and ribosome biogenesis factors (Fig. 2a), together with the nucleoid-proximal localisation (Extended Data Fig. 1a,b), suggested that CsdA condensates form a ribosome biogenesis compartment at sites of rRNA synthesis. To test this, we visualised CsdA-mEGFP together with one of the seven *E. coli* ribosomal RNA operons, the *rrnD* locus, fluorescently marked using a mCherry-ParB/parS reporter system^35^. CsdA condensates localised in close proximity to *rrnD* foci (Fig. 2b,c). Since only one of the seven *rrn* operons was fluorescently labelled, complete overlap was not expected. The spatial association between CsdA condensates and ParB/*parS* foci was lost when *rrnD* was deleted (Fig. 2c), and CsdA formed significantly fewer condensates per cell in the strains lacking *rrnD* (Extended Data Figs. 3a-c). Furthermore, transcription inhibition with rifampicin completely abolished CsdA condensates, indicating that their maintenance depends on interactions with the transcription machinery or the nascent rRNA. Likewise, CsdA condensates but not ParB/*parS* foci were disrupted by treatment with 1,6-hexanediol (Fig. 2d,e and Extended Data Figs. 1d,e, and 3a). Since both treatments also altered nucleoid morphology^36,37^, chromosome organisation may additionally contribute to foci integrity (Fig. 2d and Extended Data Fig. 3a).

We examined how other components of the transcription and ribosome biogenesis machinery are organised during cold adaptation. RNA polymerase subunit β’ (RpoC) and the rRNA anti-termination factor NusB were previously reported to colocalise within condensates during rapid growth^11,13^. Interestingly, both factors, whether ectopically expressed from plasmid (Fig. 2f,g and Extended Data Fig. 3d) or endogenously tagged (Extended Data Fig. 3e-g), assembled into prominent foci during cold adaptation that colocalised with CsdA condensates. By contrast, ObgE, a GTPase acting during late stages of ribosome assembly, remained largely diffuse. Likewise, the ribosomal proteins L1 and S2, components of the mature 50S and 30S ribosomal subunits, respectively, exhibited a diffuse cellular distribution without enrichment in CsdA condensates (Extended Data Fig. 3e,f). This localisation pattern suggested that CsdA condensates selectively recruit components of the early ribosome biogenesis machinery but not mature ribosomes.

To test whether CsdA condensates selectively recruit proteins identified in the CsdA interaction landscape, we incubated reconstituted CsdA condensates with a selection of purified candidate interactors (ObgE and HrpA) and the four other *E. coli* DDXs (SrmB, DbpA, RhlE and RhlB) (Fig. 2h,i and Extended Data Figs. 3h). SrmB, RhlE, and HrpA were efficiently enriched, whereas ObgE, DbpA and the BR-body DDX RhlB showed substantially weaker correlation, consistent with the selectivity observed by AP–MS (Fig. 2a). Thus, CsdA condensates selectively enrich a subset of RNA-associated factors rather than indiscriminately concentrating RNA-binding proteins.

In summary, these findings identify CsdA condensates as compositionally selective ribosome biogenesis compartments that spatially couple rRNA synthesis with RNA remodelling.

### The C-terminal condensation module promotes CsdA compartment assembly

To dissect the contribution of condensation to CsdA function, we sought separation-of-function mutants that impair condensate formation while retaining ATPase activity. CsdA consists of a conserved RecA ATPase core followed by an extended C-terminal region containing a dimerisation domain (DD), an RNA-binding domain (RBD), and two intrinsically disordered regions (IDRs)^38,39^ (Fig. 3a). Both IDRs are highly charged low-complexity sequences enriched in arginine and disorder-promoting residues (glycine, proline) (Extended Data Fig. 4a), features characteristic of condensate-promoting sequences (Extended Data Fig. 4b), pointing to the C-terminal region of CsdA, especially the two IDRs, being important for condensation.

**Fig. 3.**
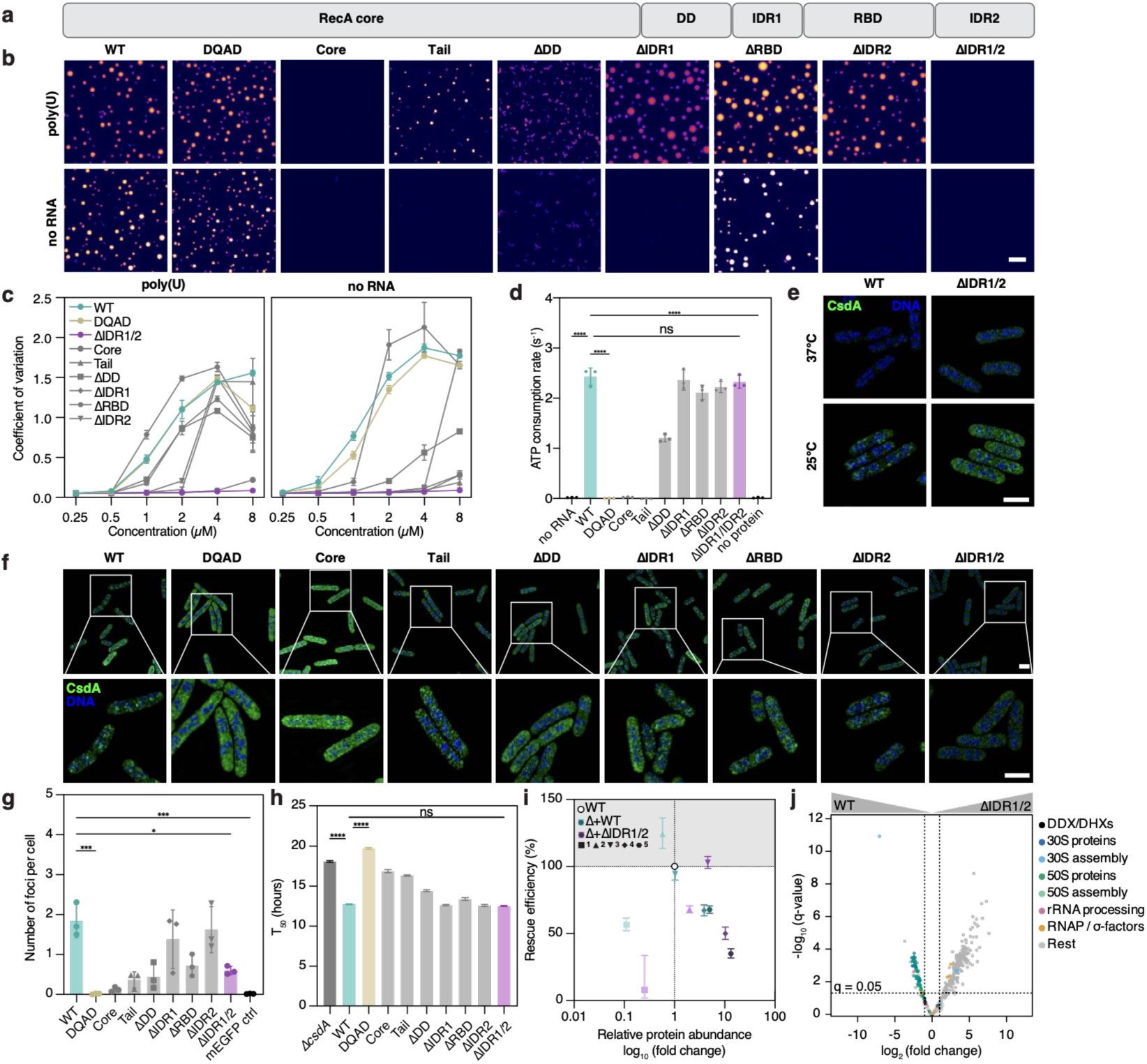
Intrinsically disordered regions promote CsdA condensation and ribosome biogenesis factor recruitment. **a,** Domain architecture of *E. coli* CsdA showing the RecA-like ATPase core, dimerisation domain (DD), RNA-binding domain (RBD), intrinsically disordered region 1 and 2 (IDR1 and IDR2). **b,** Representative in vitro condensation of 4 μM CsdA^WT^ and indicated CsdA variants ± poly(U) at 25°C (N=3). Scale bar 20 µm. **c,** Concentration-dependent quantification condensation from 3b and Extended Data Fig. 4d. **d,** ATPase activity of CsdA^WT^ and variants (N=3). **e,** 3D-SIM comparing CsdA^WT^ and CsdA^ΔIDR1/2^ at 37°C and 25°C. DNA stained with Hoechst (N=3). For all 3D-SIM images scale bar 2 µm. **f,g,** Localisation and foci per cell of endogenously mEGFP-tagged CsdA variants at 25°C (N=3). **h,** Time to reach half-max-OD600 (T_50_) during growth at 25 °C of endogenous CsdA mutants (N=3). **i,** Relationship between cellular abundance and growth-rescue of CsdA^WT^ and condensation-deficient CsdA^ΔIDR1/2^ expressed at increasing levels (1 to 5) in *ΔcsdA* cells; grey shading: conditions rescuing cold adaptation, dashed-lines: CsdA^WT^ protein abundance (N=3). **j,** AP-MS of CsdA^WT^ and CsdA^ΔIDR1/2^ interactomes; dashed lines: q=0.05 and |log_2_(FC)|=1 threshold. **c,d,g,h,i,j,** Mean ± SD. Significance assessed by ordinary one-way ANOVA.

Systematic domain dissection confirmed this prediction (Fig. 3b and Extended Data Fig. 4c). The isolated C-terminal tail formed condensates in the presence of poly(U) RNA, indicating that it contains features sufficient to support RNA-mediated multivalent interactions. Inclusion of the RNA-binding ATPase core further enhanced condensation, both for wild-type CsdA and the enzymatically inactive Walker B DQAD mutant (Fig. 3b,c and Extended Data Fig. 4c,d. and 5a). Deletion of either IDR reduced condensation but did not abolish it in the presence of RNA, whereas the ΔIDR1/2 variant remained condensation deficient, indicating that the two IDRs contribute redundantly to RNA-mediated condensation.

Next, we tested whether condensation and RNA-dependent ATP hydrolysis are mechanistically linked. As expected, CsdA^DQAD^ showed no detectable ATPase activity compared to the wild-type CsdA (Fig. 3d and Extended Data Fig. 5b), whereas condensation-deficient CsdA^ΔIDR1/2^ retained full ATPase activity. This establishes CsdA^DQAD^ and CsdA^ΔIDR1/2^ as complementary separation-of-function mutants that selectively disrupt catalytic function or condensation, respectively.

The CsdA^ΔIDR1/2^ separation-of-function variant allowed us to test how condensation contributes to CsdA foci formation in cells during cold adaptation. When expressed from the endogenous locus, CsdA^ΔIDR1/2^ formed markedly fewer nucleoid-associated compartments than wild-type CsdA (Fig. 3e-g, and Extended Data Fig. 5c,d). ATPase- and RNA-binding-deficient variants likewise impaired foci formation (Fig. 3f,g, and Extended Data Fig. 5d), indicating that efficient cellular compartment formation requires coordinated contributions from the C-terminal condensation module, RNA binding, and ATP turnover.

Since CsdA condensation is concentration-dependent (Fig. 1d,e), we assessed its contribution to growth using a titratable expression system. Wild-type CsdA or CsdA^ΔIDR1/2^ were expressed from promoters of graded strength (Extended Data Fig. 6a-d) in a *ΔcsdA* background to determine the expression levels required to rescue the severe low-temperature growth defect caused by loss of CsdA at 25 °C (Fig. 3h, and Extended Data Fig. 6e-i). Protein abundance was quantified by western blot, normalised to the endogenous protein expression levels and related to rescue efficiency of the cold adaptation (Fig. 3i, and Extended Data Fig. 6j-l). Compared to wild-type CsdA, CsdA^ΔIDR1/2^ required higher relative expression levels to achieve comparable rescue (Fig. 3i). This is consistent with the observation that endogenous CsdA^ΔIDR1/2^ accumulated to significantly higher levels than wild-type CsdA and caused no significant growth defect at 25 °C (Fig. 3h, and Extended Data Fig. 1f-i, 6e-i). The increased abundance may compensate for its reduced ability to achieve high local concentrations through condensation. Interestingly, excessive CsdA expression reduced rescue efficiency, indicating that CsdA abundance must be balanced as both insufficient and excessive levels impair fitness during cold adaptation.

The reduced condensation capacity of CsdA^ΔIDR1/2^ was also reflected in its interaction landscape. AP-MS of CsdA^ΔIDR1/2^ enriched ribosome biogenesis factors, but to a significantly lower extent than wild-type CsdA (Fig. 3j, and Extended Data Fig. 6o), consistent with impaired assembly of the broader CsdA compartment. Because the deleted IDRs likely mediate protein and RNA interactions, the reduced recovery of ribosome biogenesis factors can however not be attributed exclusively to defective condensation.

In summary, we identified the C-terminal IDRs as key determinants of CsdA condensation in vitro and compartment formation in cells. Condensation enhances the rescue efficiency of CsdA during cold adaptation. Increased protein abundance can partially compensate for impaired condensation, but tight regulation of CsdA abundance is important for cellular fitness.

### CsdA compartmentalisation promotes ribosome assembly under cold conditions

How does condensation enhance CsdA function during *E. coli* cold adaptation? Given the close association of CsdA condensates with sites of ribosome biogenesis, we asked whether disrupting CsdA compartment formation impairs ribosome maturation, especially during cold adaptation. Ribosome profiles were analysed in strains lacking CsdA (*ΔcsdA*) or expressing condensation-deficient CsdA^ΔIDR1/2^, ATPase-deficient CsdA^DQAD^, or CsdA^WT^ from the endogenous locus. At 37 °C, ribosome maturation was largely unaffected in all strains (Fig. 4a). At 25 °C, strains lacking CsdA^40–42^ or expressing the CsdA^DQAD^ variant displayed severe ribosome maturation defects, including reduced mature subunit peaks and accumulation of aberrant pre-50S intermediates (Fig. 4b,c and Extended Data Fig. 7a). CsdA^ΔIDR1/2^ produced a similar phenotype like *ΔcsdA* and DQAD, with accumulation of the pre-50S intermediates, but milder effects on the 30S peak. Individual deletion of IDR1 or IDR2 caused milder defects for both subunit peaks (Extended Data Fig. 7b,c). These findings demonstrate that both ATP hydrolysis and the C-terminal condensation module contribute substantially to CsdA function in ribosome biogenesis during cold adaptation.

**Fig. 4.**
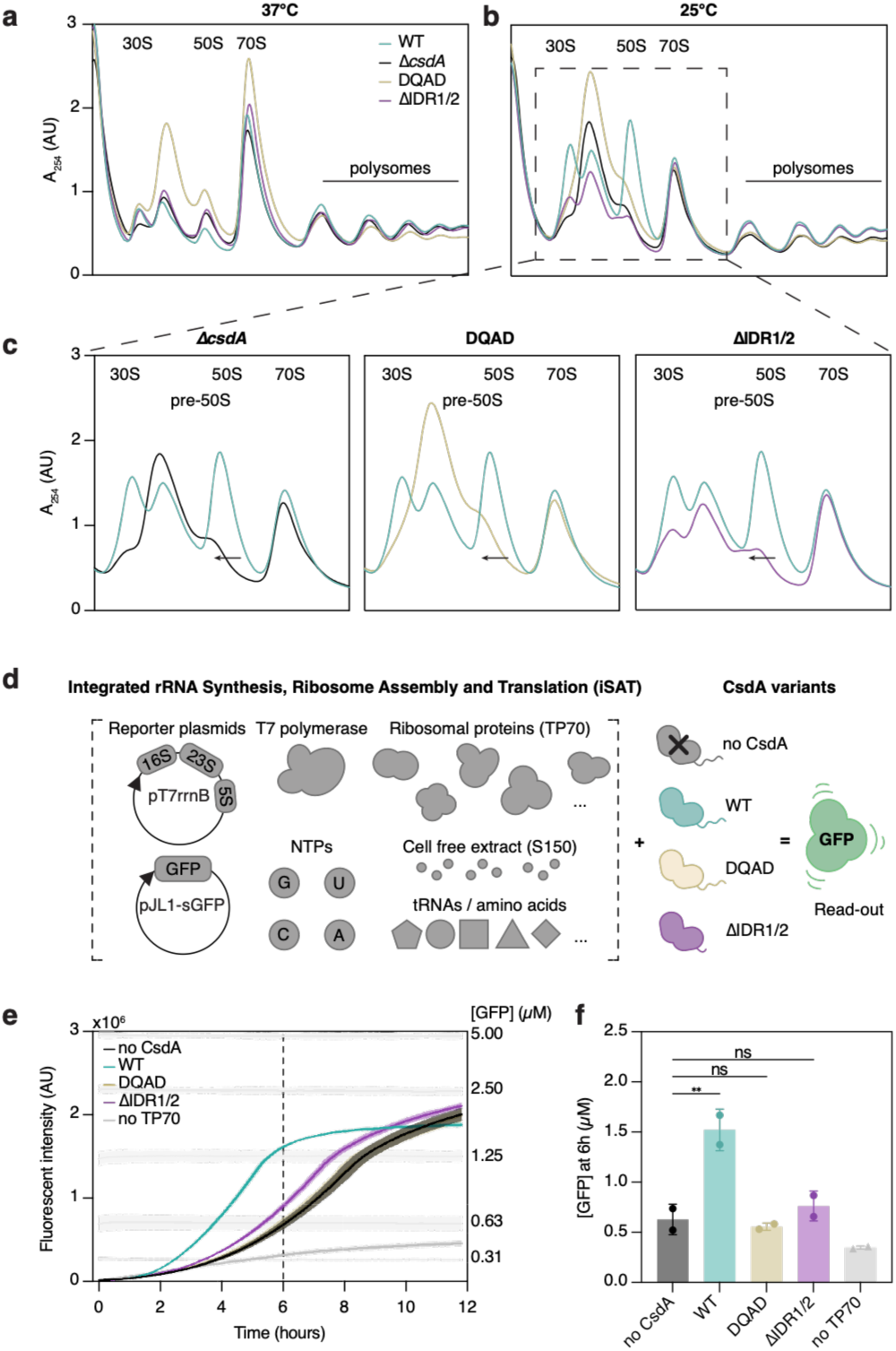
CsdA condensation promotes ribosome biogenesis. **a,b,** Representative sucrose-gradient profiles of cells expressing wild-type CsdA, CsdA^ΔIDR1/2^, CsdA^DQAD^ or lacking CsdA (*ΔcsdA*) grown at 37°C or 25°C; positions of 30S, pre-50S, 50S, 70S and polysome fractions are indicated (N=3). **c,** Enlarged 25°C profiles highlighting altered pre-ribosomal populations; the black arrow marks the shift from the 50S to pre-50S intermediate peak. **d,** Schematic of the integrated synthesis, assembly and translation (iSAT) assay used to monitor de novo ribosome biogenesis. Reactions contained 16S, 23S and 5S rRNA and sfGFP templates, TP70 ribosomal proteins, tRNAs, amino acids, NTPs, S150 extract and an ATP-regeneration system. Recombinant wild-type CsdA, CsdA^ΔIDR1/2^, CsdA^DQAD^ or no CsdA was added as indicated. Translation is read-out by GFP reporter accumulation. **e,** Representative GFP reporter accumulation over time during iSAT reactions at 25°C ± indicated CsdA variants and standard GFP fluorescence. Solid lines indicate mean of technical triplicates ± SD. The dashed-line indicates the 6 h time-point used for GFP quantification (N=2). **f,** Quantified GFP concentration after 6 h iSAT at 25°C (N=2). Mean ± SD shown as bar, individual replicates shown as dots. Significance assessed by ordinary one-way ANOVA.

To determine whether CsdA condensation directly promotes ribosome assembly, we turned to a reconstituted in vitro ribosome assembly system (iSAT)^43^, in which ribosomes co-transcriptionally assemble *de novo* from isolated ribosomal proteins and rRNA actively transcribed from a plasmid encoding the complete *rrnB* operon in the presence of crude S150 *E. coli* cell extract (Fig. 4d). Ribosome assembly and function was monitored by translation of a GFP reporter. Lowering the reaction temperature from 37 °C to 25 °C markedly reduced ribosome assembly efficiency, consistent with the idea that low temperature imposes RNA-folding barriers, but only minimally affected translation efficiency of mature, pre-assembled 70S ribosomes (Extended Data Fig. 7d-g). Addition of 1.25 µM CsdA^WT^, a concentration at which CsdA robustly condensed in iSAT reaction buffers (Extended Data Fig. 7h), accelerated GFP reporter accumulation (Fig. 4e,f and Extended Data Fig. 7i-k). By contrast, condensation-deficient CsdA^ΔIDR1/2^ or ATPase-deficient CsdA^DQAD^ failed to significantly enhance reporter accumulation.

Together, these findings demonstrate that in addition to its ATPase activity, CsdA condensation promotes ribosome assembly under cold conditions. This identifies CsdA condensation as an adaptive mechanism that spatially concentrates RNA remodelling activity to facilitate efficient ribosome biogenesis.

### Convergent evolution of CsdA-mediated condensation across bacteria

How widely distributed is this mechanism for cold adaptation in bacteria? Comparative genomics showed that the CsdA gene family is broadly distributed across the bacterial tree (121/290 genomes spanning 12 of 14 lineages examined; Extended Data Fig. 8a,b). Bioinformatic analysis of the C-terminal module of CsdA orthologs indicates that condensation propensity has evolved multiple times independently, particularly in *E. coli* and its gamma-proteobacterial relatives (Extended Data Fig. 8b,c). Consistent with our experimental analyses, these evolutionary changes preferentially map to IDR2, identifying this region as the principal condensation module. By contrast, the C-terminal module has been lost entirely in some thermophilic lineages, with a significant phylogenetically controlled correlation between the presence of IDR2 and optimal growth temperature (MWU p = 0.0015, phyloglm p = 0.02; Extended Data Fig. 8d-e). A similar trend was observed in the smaller set of directly measured growth temperatures (number of genomes = 25, TEMPURA) (Extended Data Fig. 8d). Taken together with our experimental findings, this suggests that eukaryotes, and various lineages of bacteria, have convergently evolved a common, condensation-mediated mechanism to promote ribosome biogenesis.

## Discussion

Our work demonstrates that spatial compartmentalisation of ribosome biogenesis is not unique to eukaryotes. During cold adaptation, *E. coli* transiently assembles a compartment that spatially couples rRNA synthesis, RNA remodelling and ribosome assembly, thereby sustaining efficient ribosome biogenesis when RNA folding becomes limiting.

The adaptive nature of this compartment emerges from concentration-dependent regulation of CsdA condensation: maintaining CsdA close to its condensation threshold during optimal growth allows a relatively modest increase in protein abundance to trigger compartment assembly while avoiding the fitness cost associated with constitutively high CsdA expression. This threshold-dependent regulation concentrates RNA-remodelling activity precisely when and where it is needed.

Condensation thus enhances ribosome biogenesis not by introducing a new biochemical activity, but by spatially organising an existing one. Concentrating CsdA at sites of rRNA synthesis promotes efficient remodelling of the RNA-RNA interactions that accumulate in such RNA-rich environments, while limiting non-productive encounters with off-target RNAs and unnecessary ATP hydrolysis^44–48^. Similar organisational principles operate in the eukaryotic nucleolus, where sequential rRNA processing and DEAD-box ATPases continuously resolve RNA entanglements^21,49–51^.

Chromosome organisation may further contribute to establishing these compartments. The seven *E. coli rrn* operons have been proposed to undergo partial spatial clustering^35^, potentially creating a local scaffold of nascent rRNA that nucleates CsdA condensation. Such a mechanism could explain the sensitivity of CsdA condensates to perturbations of transcription or nucleoid organisation^36,37^.

Previous work established that during rapid bacterial growth, clusters of RNA polymerase and antitermination factors assemble transcription factories at highly transcribed *rrn* operons, where they enhance rRNA synthesis and have been proposed to support early co-transcriptional rRNA folding^7,13,14,52–54^. Our findings extend this organisational principle beyond transcription by showing that cold adaptation recruits downstream RNA-remodelling and ribosome assembly activities into a specialised compartment. Rather than organising a single stage of ribosome production, the CsdA compartment therefore integrates successive stages of ribosome biogenesis.

This organisation bears striking parallels to the eukaryotic nucleolus, whose multilayered architecture coordinates successive stages of rRNA processing and pre-ribosomal maturation^2,49,50,55^. The bacterial CsdA compartment is likely considerably simpler and may not encompass the entire ribosome biogenesis pathway. ObgE, L1 and S2 – all relatively late or peripheral factors – remained predominantly diffuse, suggesting that CsdA compartments preferentially support early and co-transcriptional stages of ribosome assembly^5,56^, before more mature pre-ribosomal particles leave the compartment^42^.

A major distinction, however, is the conditional nature of the bacterial compartment. Whereas eukaryotes rely on permanent spatial organisation to sustain their elaborate ribosome biogenesis pathway, CsdA condensates emerge specifically during cold adaptation, when stabilised RNA structures increase the demand for rRNA remodelling and make ribosome assembly thermodynamically and kinetically challenging. This adaptive organisation allows bacteria to spatially reorganise ribosome biogenesis only when it becomes advantageous, and to return to a diffuse state when those constraints are relieved. The widespread occurrence of condensation-prone CsdA/CshA-related DDXs suggests that this strategy extends well beyond *E. coli* and that similar compartments may play a more persistent role in psychrophilic bacteria, where ribosome biogenesis continuously operates at temperatures that favour stable RNA structures^57^.

More broadly, our findings suggest that biomolecular condensates can emerge as adaptive modules when environmental conditions challenge the efficiency of essential biochemical processes. Rather than requiring new enzymatic activities, environmental adaptation arises through spatial reorganising of existing biochemical activities. Facultative compartments of this kind may provide a simple solution towards biochemical robustness and, over evolutionary time, could represent intermediates from which more constitutive cellular organisation emerges. In this view, the CsdA compartment represents a convergent route towards nucleolus-like organisation.

## MATERIAL AND METHODS

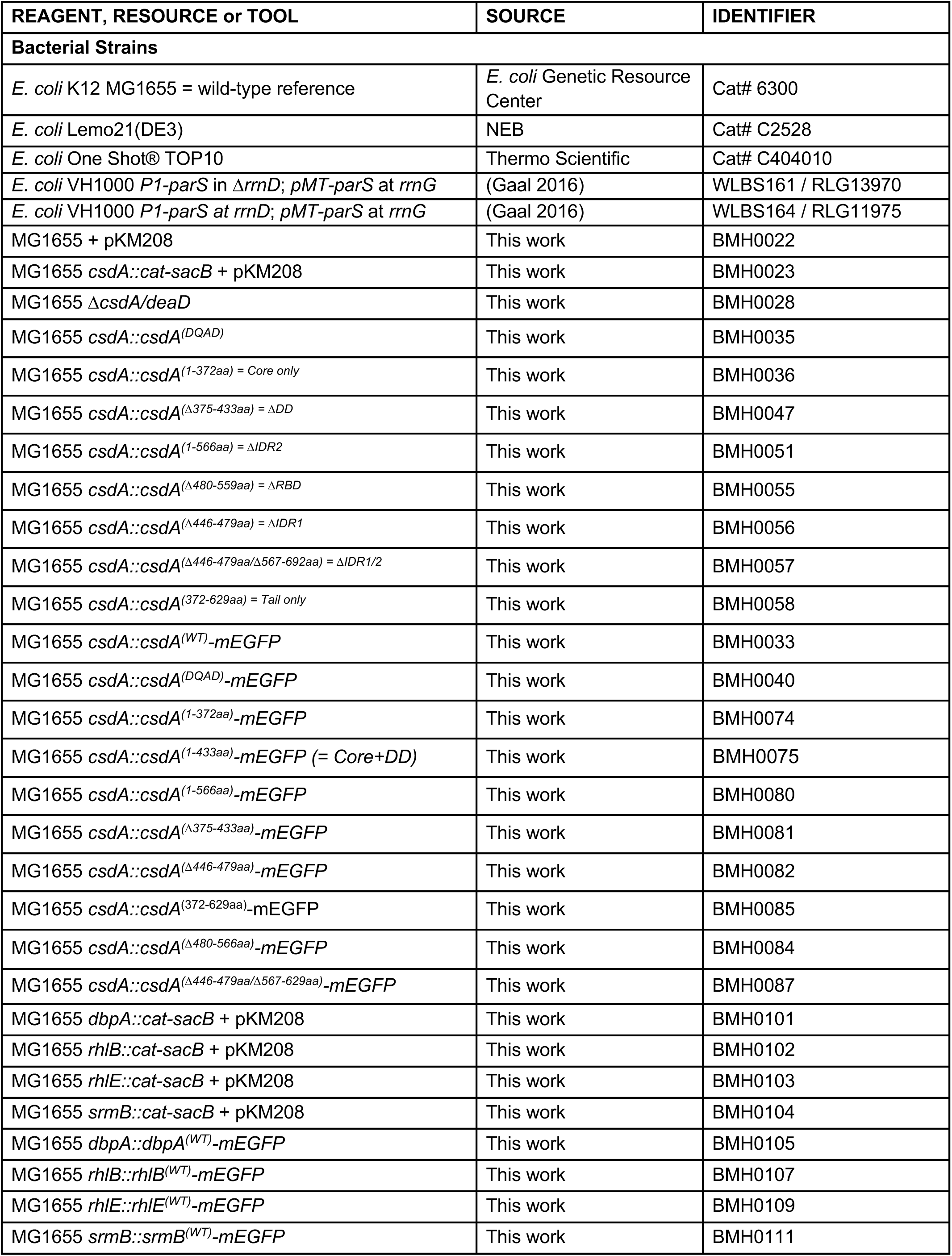
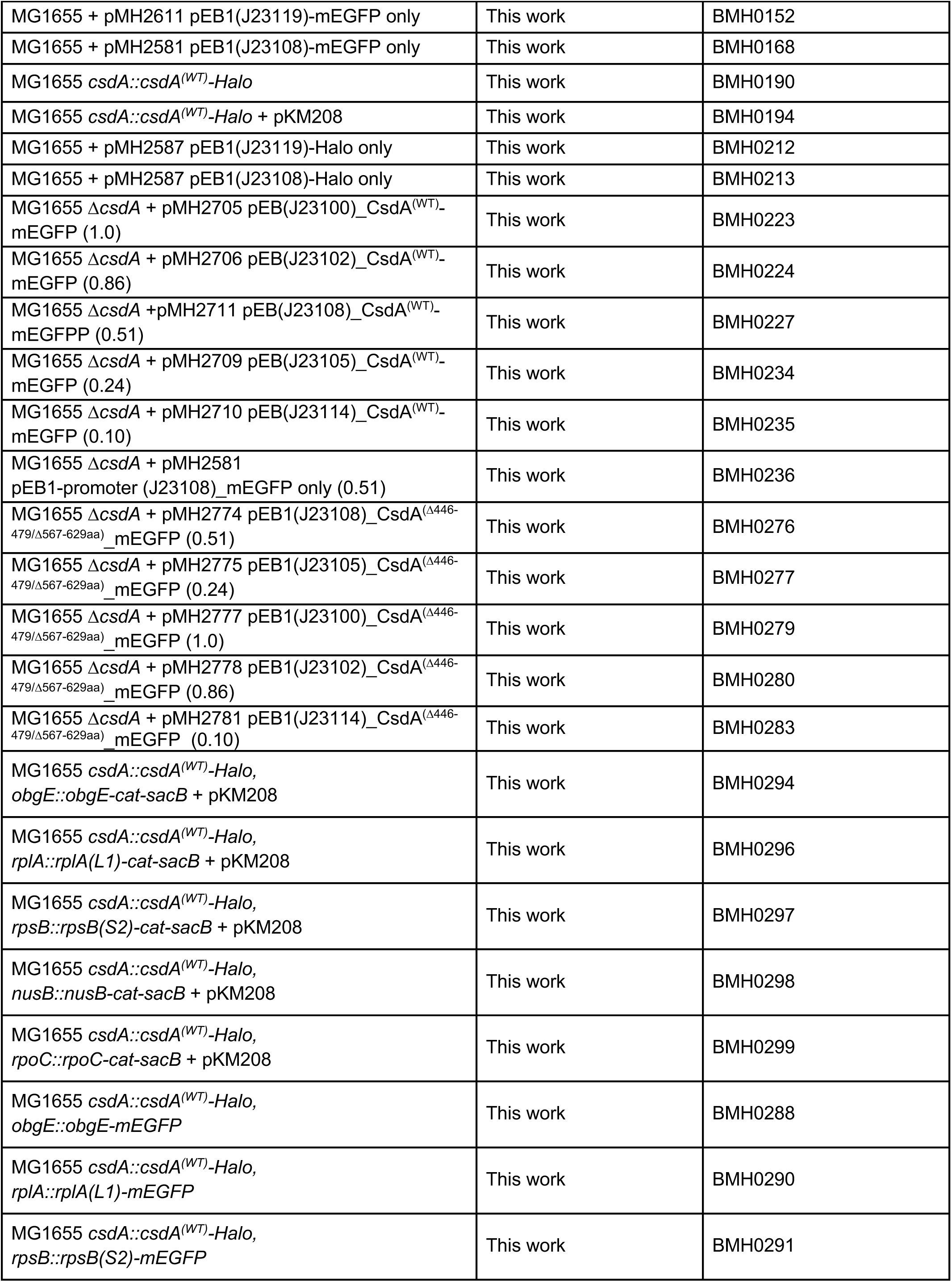
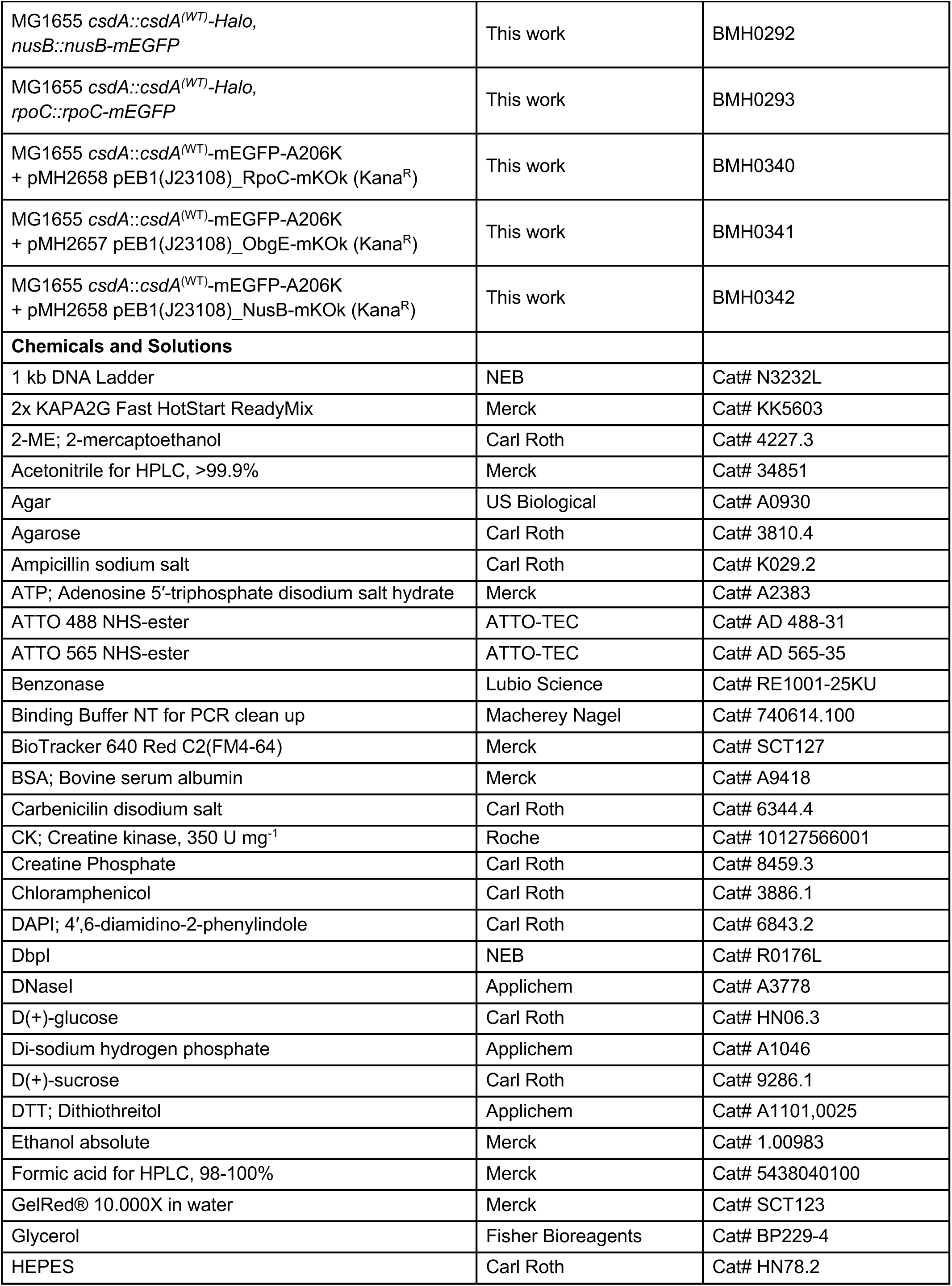
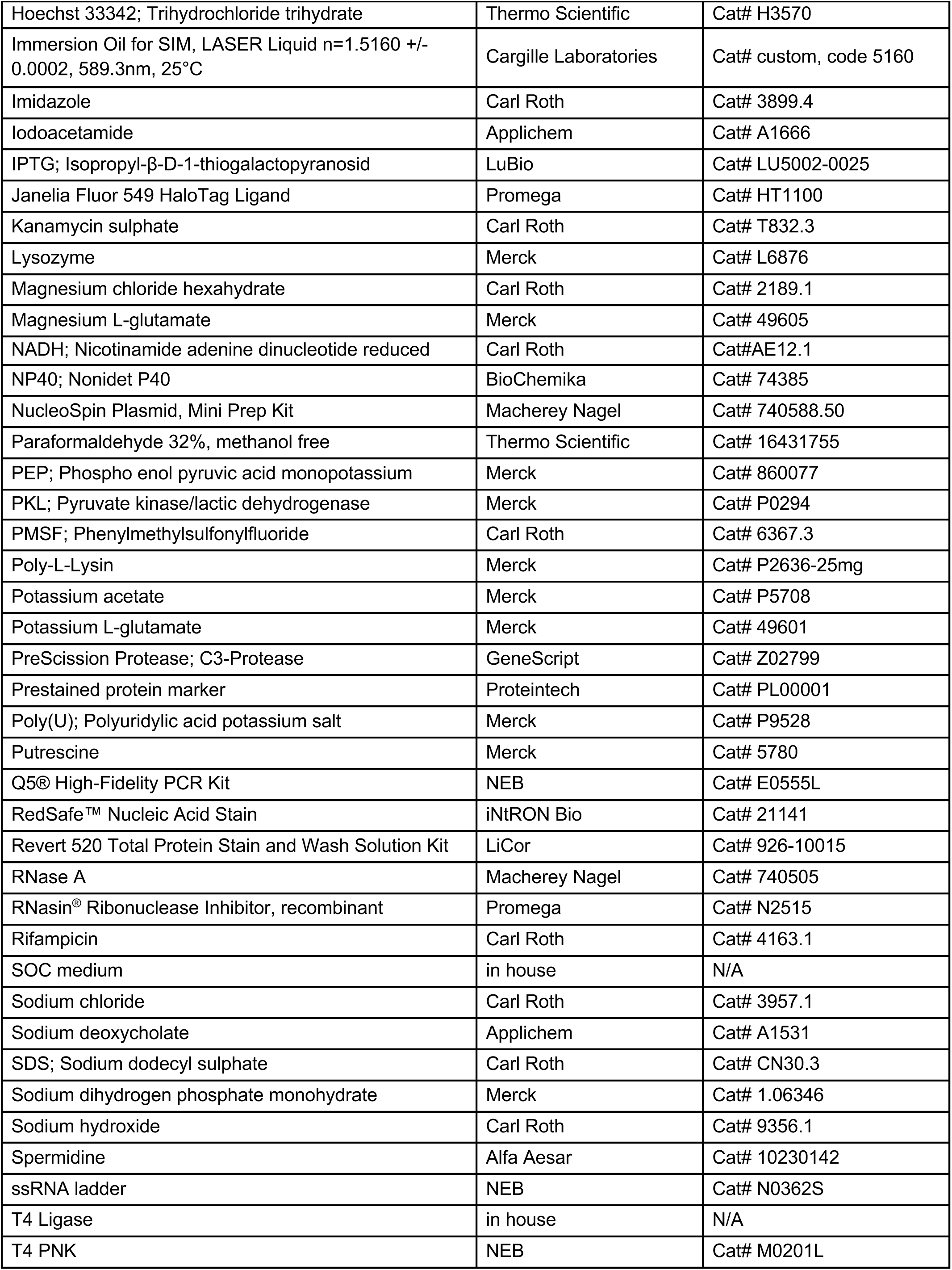
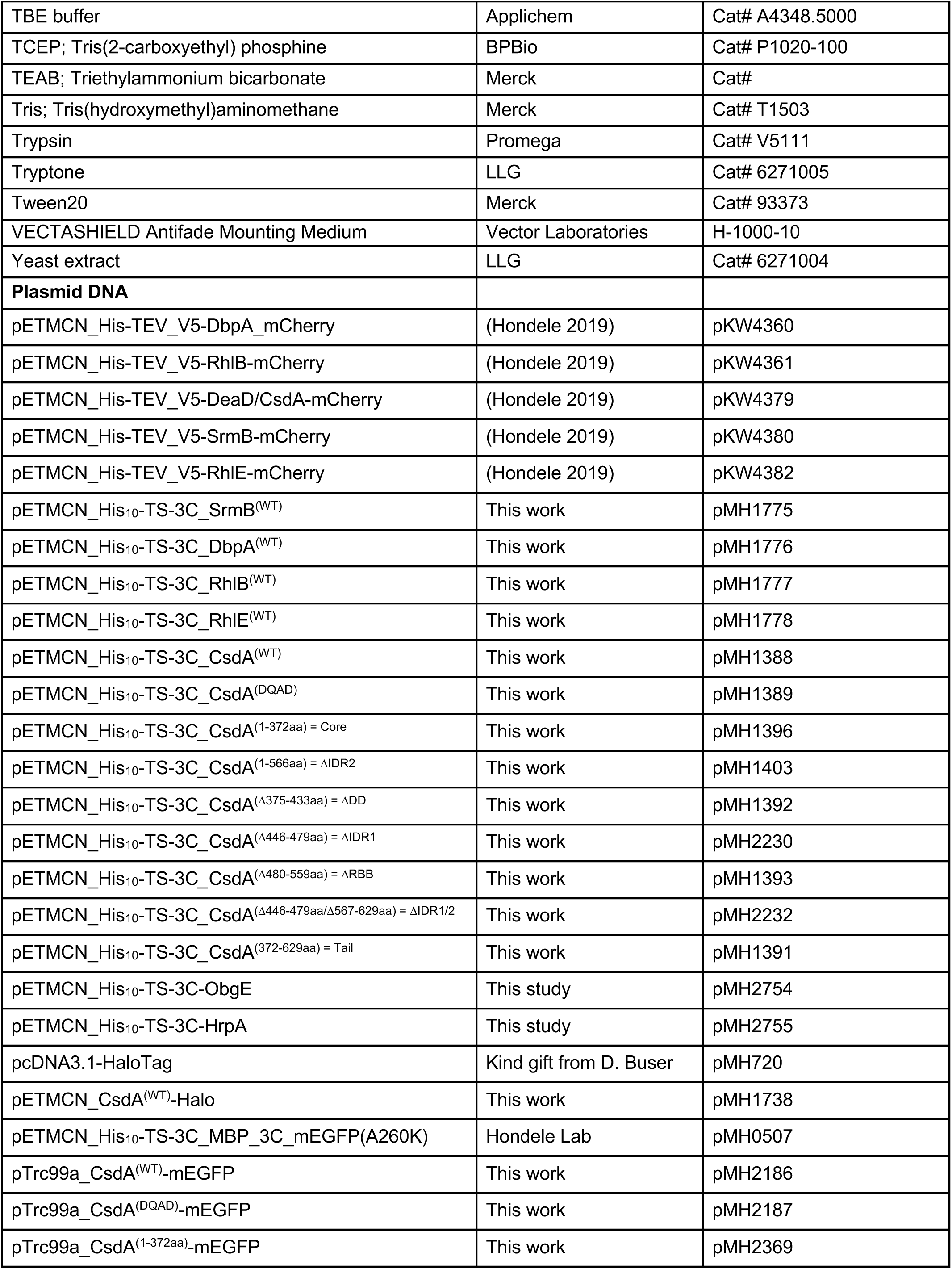
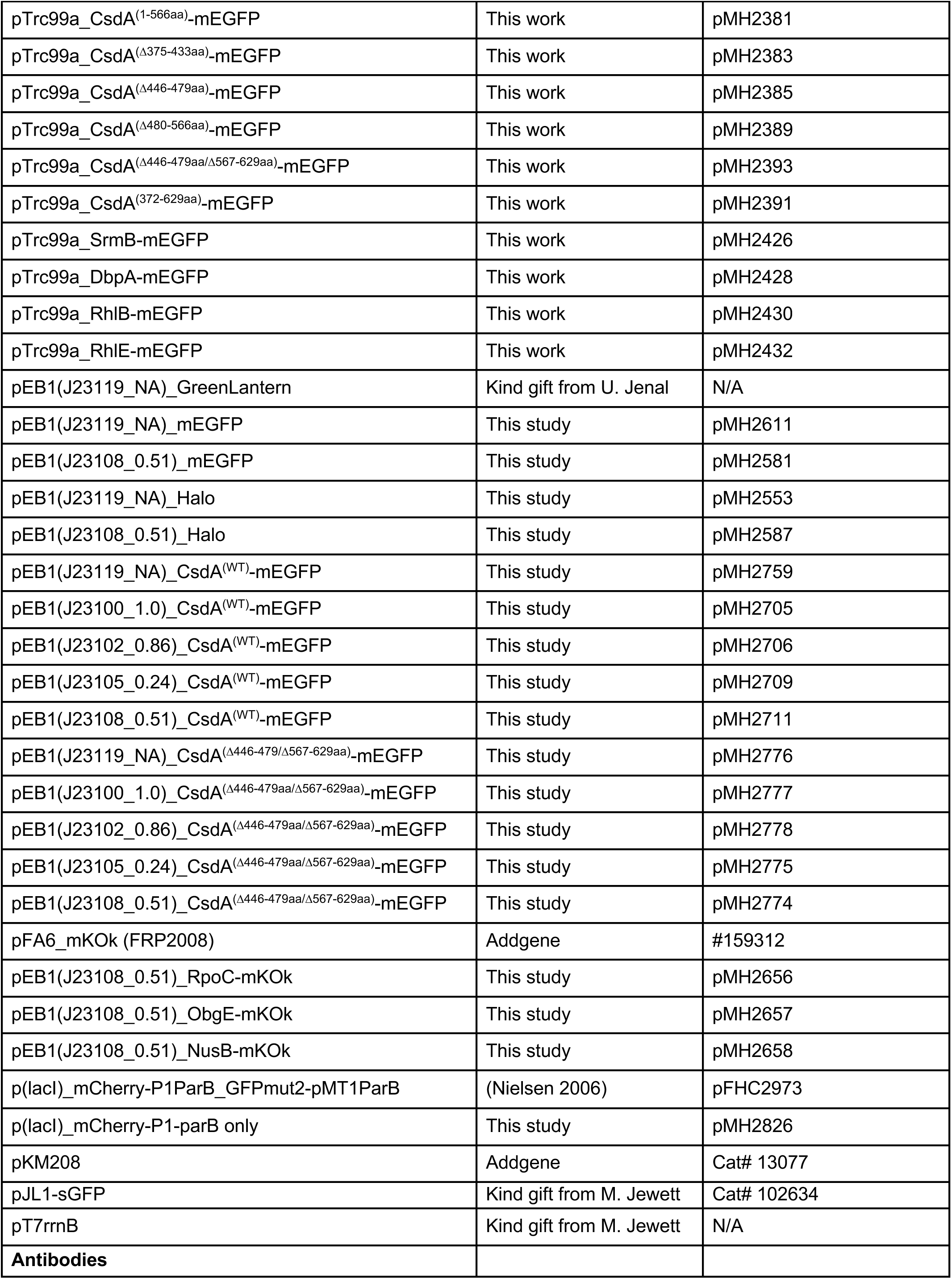
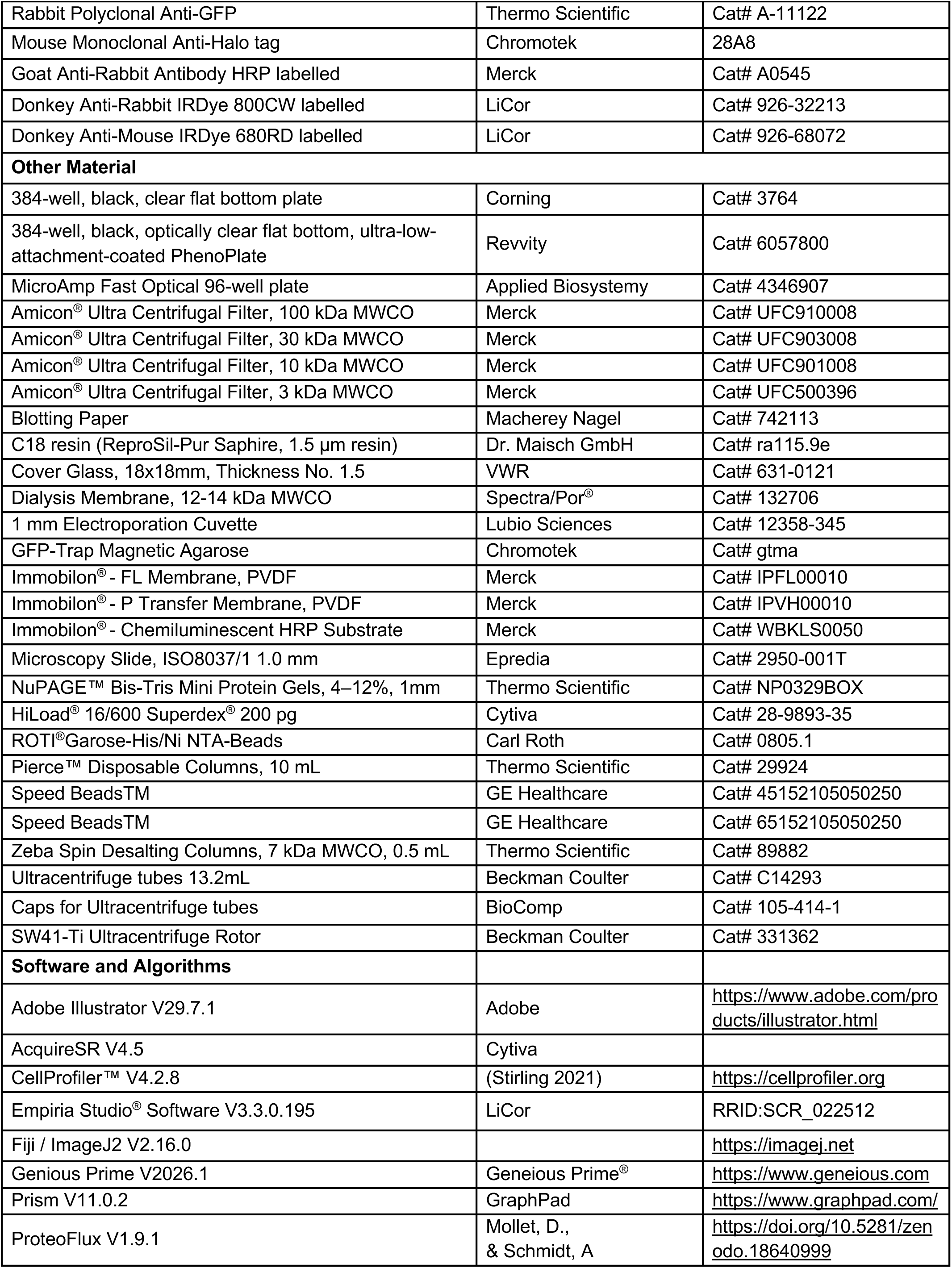
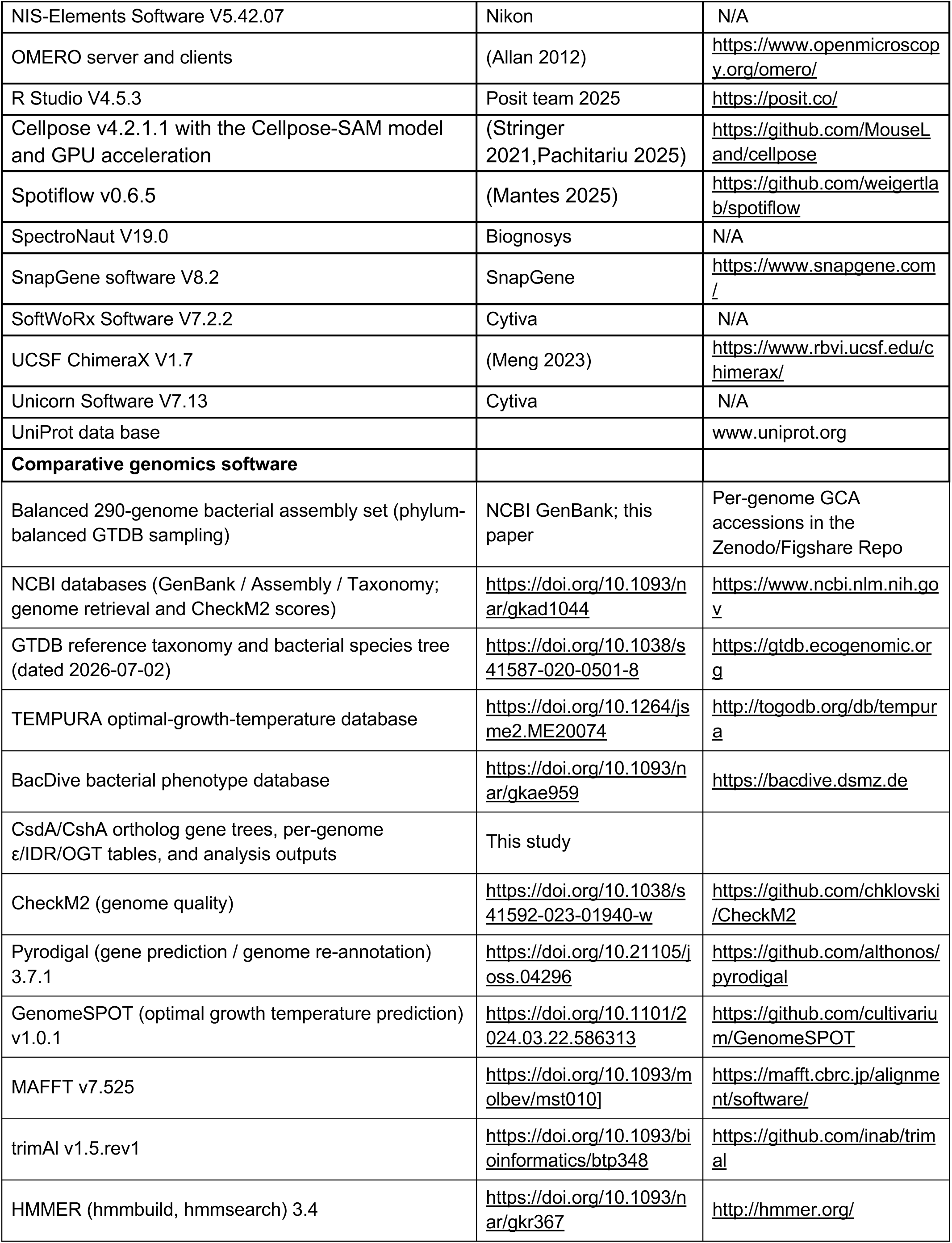
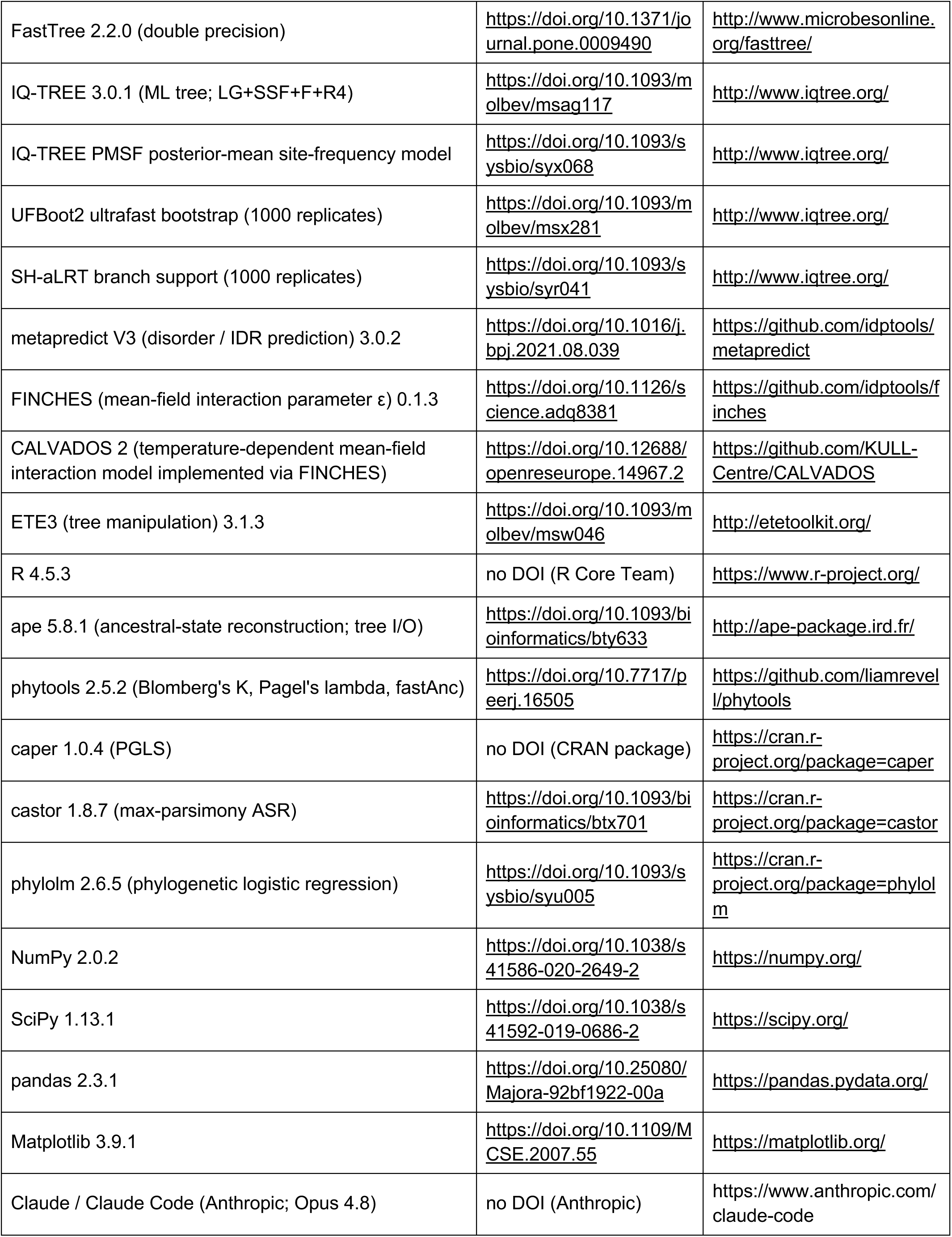
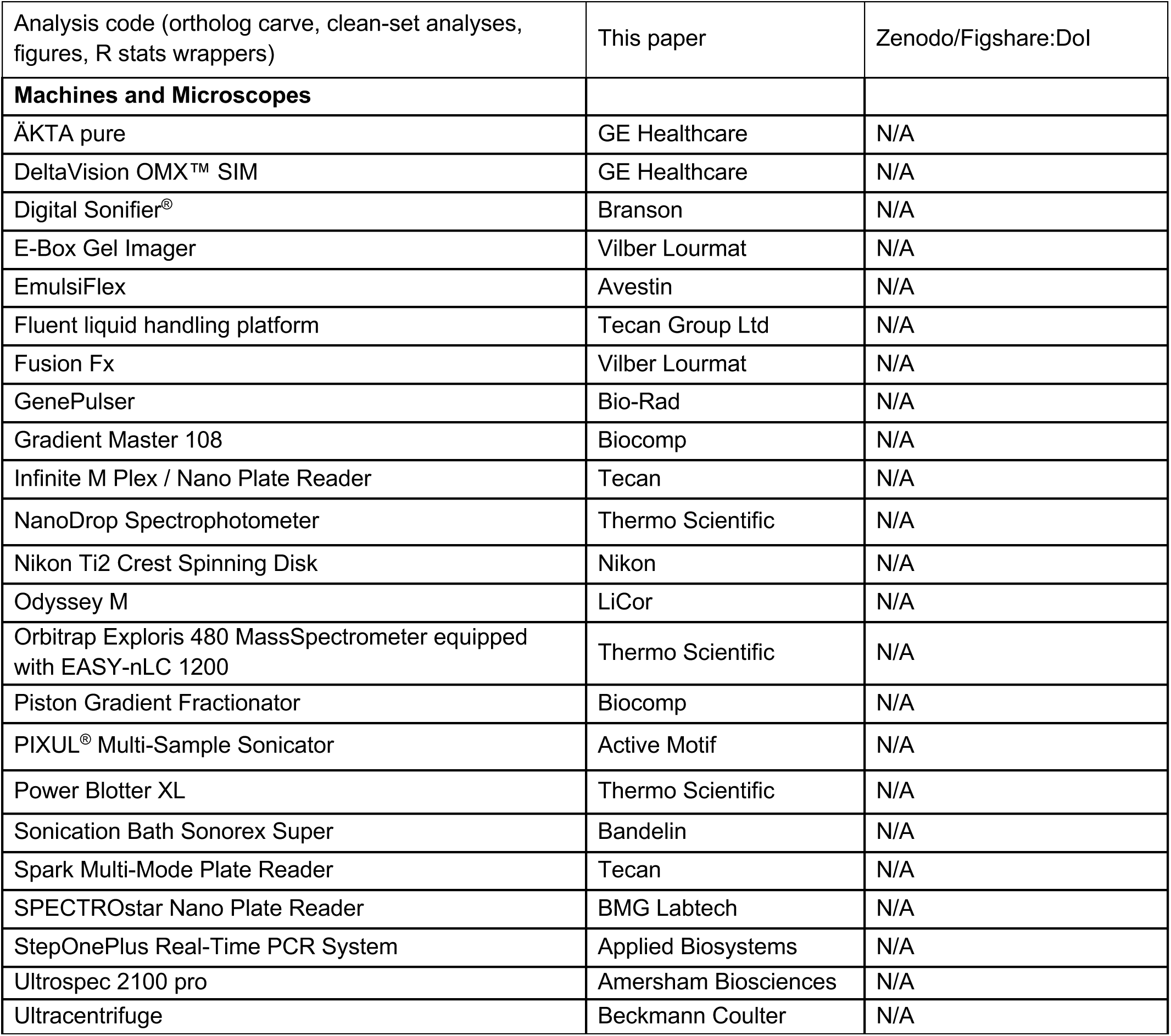

### Bacterial strains and growth conditions

Bacterial strains used in this study are listed in (Key Resources Table). *E. coli* Top10 cells were used for molecular cloning and plasmid propagation, while *E. coli* Lemo21(DE3) cells were used for recombinant protein expression. *E. coli* K-12 MG1655 was used as the wild-type reference strain and as the parental background for generation of chromosomally tagged and mutant strains using the λ-Red recombination system.

Bacterial strains were stored as glycerol stocks at −75°C. For strain propagation, glycerol stocks were streaked onto lysogeny broth (LB) agar plates supplemented with the appropriate antibiotics and incubated at 37°C unless otherwise stated. Liquid cultures were grown in LB medium at the indicated temperatures (37°C, 30°C, or 25°C) with shaking at 100–200 rpm. Selection antibiotics were added when required at the following final concentrations: kanamycin (50 μg mL⁻¹), ampicillin (100 μg mL⁻¹), carbenicillin (100 μg mL⁻¹), chloramphenicol (30 μg mL⁻¹), or D-sucrose (8%).

### Plasmid construction

All parental plasmids, newly generated constructs, cloning strategies and oligonucleotides used in this study are listed in (Key Resources Table). Unless otherwise indicated, DNA fragments were amplified using Q5 High-Fidelity DNA Polymerase (New England Biolabs) and assembled by Gibson assembly, restriction enzyme-based cloning or Q5-based mutagenesis. Coding sequences were amplified either from *E. coli* K-12 MG1655 genomic DNA or from pre-existing plasmid templates.

For recombinant protein expression and purification, genes encoding CsdA, SrmB, DbpA, RhlB, RhlE, HrpA and ObgE were cloned into ampicillin-resistant, IPTG-inducible pETMCN expression vectors. The resulting constructs encoded an N-terminal His_10_-TwinStrep affinity tag followed by a 3C protease cleavage site and the protein of interest, allowing removal of the affinity tag after purification. The five *E. coli* DDX genes were either subcloned from previously generated pETMCN constructs^1^ or amplified from *E. coli* genomic DNA.

CsdA mutant and truncation variants were generated from the pMH1388 pETMCN-His_10_-TwinStrep-3C-CsdA^(WT)^ construct using Q5-based site-directed mutagenesis. Internal deletions and truncations were introduced by selective amplification of the desired plasmid regions followed by phosphorylation and ligation of the linearized PCR products, whereas amino-acid substitutions, including the ATPase-deficient CsdA^DQAD^ variant, were introduced using mutagenic primers. Boundaries of the CsdA truncation constructs were selected based on the previously described domain architecture of CsdA^2^ together with sequence-based predictions of regions contributing to condensation. Liquid-liquid phase-separation propensity was assessed using the FuzDrop algorithm^3^, using either the corresponding UniProt accession number or the amino-acid sequence of the respective CsdA variant. Protein sequences, UniProt accession numbers and construct boundaries are provided in Extended Data Table 2.

For generation of templates used for chromosomal fluorescent tagging by λ-Red recombination, coding sequences were cloned into pETMCN- or pTrc99a-derived plasmids containing either mEGFP or HaloTag. The fluorescent or HaloTag coding sequence was positioned C-terminally to the protein of interest and separated from the coding sequence by an in-frame flexible GSGGSGG linker. The mEGFP sequence used for cloning was amplified from plasmid pMH507 (pETMCN_His_10_-TS-3C_MBP_3C_mEGFP(A260K)), mKOk from FRP2008 (Addgene #159312) the HaloTag from pcDNA3.1-HaloTag (a kind gift from D. Buser). These constructs subsequently served as templates for amplification of the corresponding tagging cassettes.

For constitutive expression experiments, plasmids were derived from the pEB1(PJ23119)-mGreenLantern vector obtained from the Jenal laboratory. The mGreenLantern coding sequence was replaced by either mEGFP(A206K), mKOk or HaloTag, and the parental J23119 promoter was subsequently exchanged for promoters of defined relative strengths from the Anderson constitutive promoter collection (https://parts.igem.org/Promoters/Catalog/Anderson; Extended Data Table 1). Promoters ranging from BBa_J23100, with a relative activity of 1.0, to BBa_J23114, with a relative activity of 0.10, were used to generate constructs spanning a broad range of expression levels. Coding sequences were inserted upstream of the fluorescent tag and separated by an in-frame GSGGSGG linker where applicable. This plasmid collection was used to express CsdA^WT^ and CsdA^ΔIDR1/2^ at graded levels for complementation experiments in the Δ*csdA* background, and to generate fluorescent-tag-only controls. In addition, ObgE, NusB and RpoC were expressed as fluorescently tagged constructs for colocalisation experiments with chromosomally tagged CsdA. The promoter identity and relative strength used for each construct are specified in (Extended Data Table 1).

For visualisation of *parS*-labelled *rrnD* loci, an IPTG-inducible mCherry-P1-ParB expression plasmid was generated from the previously described pFHC2973 construct encoding P1ParB-GFPmut2 and pMT1ParB^4^. The GFPmut2-pMT1ParB portion of the parental construct was removed by Q5-based mutagenesis, retaining the mCherry-P1-ParB module under control of the inducible promoter. The resulting plasmid was transformed into strains carrying *parS-*labelled *rrnD* or Δ*rrnD* loci and used to visualise chromosomal *parS* sites through recruitment of mCherry-P1-ParB.

### Transformation

Plasmids or linear DNA fragments were transformed into *E. coli* by electroporation of 20-200 ng of DNA into 10 mL of exponentially growing day cultures washed three times with sterile 10 % glycerol using a GenePulser (BioRad) with the following settings: 25 μF, 400 Ω, 1.75 kV, in 1 mm cuvettes. All steps were performed at 4°C. 1 mL Super Optimal Medium (SOC) medium was added immediately after the pulse and cells were incubated 1-4 h at 37°C (or 30°C for strains containing heat-sensitive pKM208) shaking before plating on LB plates containing the appropriate selection. Plasmids used in this study are listed in (Key Resource Table).

### Mutant construction and endogenous tagging

All strains were built in the *E. coli* K12 MG1655 background. Endogenous tagging and mutant strain generation was performed as previously described by lambda recombination using the pKM208 plasmid carrying the enzymes required for DNA recombination (Addgene, Cat# #13077) and homology flanked linear DNA fragments carrying tagged or modified genes of interest^5,6^.

In brief, lambda red expression was induced by IPTG. Electrocompetent, induced cells were transformed with PCR amplified linear DNA fragments containing 50 bp homology arm flanked *cat-sacB* selection cassettes targeting specific genomic loci. Cassette integration was tested by chloramphenicol selection and confirmed by colony PCR targeting modified genomic locus.

Strains harbouring selection cassettes and pKM208 were induced and transformed with 50 bp homology flanked PCR products carrying the genes of interest with the respective mutations and/or C-terminal protein tags. Recombination was assessed by sucrose selection, followed by a chloramphenicol counter selection.

Mutant construction or endogenous tagging was verified by colony PCR and Sanger sequencing. For double tagged *E. coli* strains an additional round of lambda recombination was performed, followed by Sanger sequenced of both modified genomic loci for verification. Expression of the tagged genes were verified by Western Blot against the integrated tag. If applicable, generated strains were further transformed with constitutive or IPTG-inducible plasmids as control or for visualisation of additional proteins.

### P1 Phage Transduction

P1vir-mediated transduction was used to introduce *parS*-labelled *rrnD* operon from donor strains into chromosomally tagged *E. coli* recipient strains expressing CsdA-mEGFP. Donor strains carrying P1-*parS(*Kana^R^) at *rrnD* and pMT-parS(Cm^R^) at *rrnG* (RLG11975) or P1-*parS*(Kana^R^) in Δ*rrnD and* pMT*-parS*(Cm^R^) at *rrnG* (RLG13970)^7^ (a kind gift from S. Weber) carrying the desired P1-*parS* insertion and antibiotic resistance marker were grown overnight in LB medium supplemented with the appropriate antibiotic.

To generate P1 lysates, overnight cultures were diluted 1:20 into 5 mL LB medium without antibiotics and supplemented with CaCl₂ and MgSO₄ to final concentrations of 10 mM and 20 mM, respectively. P1*vir* phage lysate from wild-type *E. coli* K12 MG1655 (a kind gift from U. Jenal) was added, and cultures were incubated at 37°C with shaking until complete lysis was observed (typically 2–3 h). Chloroform was then added to a final concentration of approximately 17% (v/v), samples were vortexed thoroughly, and cellular debris was removed by centrifugation at 4,000 ×g for 5 min. The phage-containing supernatant was transferred to fresh chloroform-resistant tubes, supplemented with a small amount of chloroform, and stored at 4°C. For transduction, recipient strains were grown overnight in LB medium and mixed 1:1 with fresh LB medium. CaCl₂ and MgSO₄ were added to final concentrations of 10 mM and 20 mM, respectively, followed by addition of donor-derived P1 lysate. The infection mixture was incubated for 20–30 min at 37°C without shaking to allow phage adsorption. Transduction was terminated by addition of sodium citrate to a final concentration of 100 mM, and cells were recovered in fresh LB medium for 1 h at 37°C with shaking to allow phenotypic expression of the antibiotic resistance marker. Cells were subsequently pelleted by centrifugation (4,000 ×g, 5 min) and plated on LB agar supplemented with the appropriate antibiotic and 20 mM sodium citrate to prevent further phage infection. Plates were incubated overnight at 37°C. Putative transductants were purified by repeated restreaking on selective medium until no phage plaques were observed. Correct transduction and integration of the *parS*-labelled *rrnD* or *ΔrrnD* locus was verified by colony PCR using primers flanking the insertion site, while maintenance of the chromosomal CsdA-mEGFP fusion was confirmed using primers flanking the *csdA* locus. PCR products were purified and validated by Sanger sequencing.

Strains carrying the parS-labelled *rrnD* locus, the *parS*-labelled *ΔrrnD* locus, and the corresponding parental CsdA-mEGFP strain containing the native unlabelled *rrnD* locus were transformed with ampicillin resistant plasmid pMH2826, which expresses mCherry-P1-ParB under an IPTG-inducible promoter. In strains containing a chromosomal *parS* insertion, mCherry-P1-ParB binding to *parS* enabled visualisation of the labelled *rrnD* genomic locus. The parental CsdA-mEGFP strain lacking a *parS* insertion was transformed with the same plasmid and served as a control for nonspecific or background mCherry-P1-ParB fluorescence.

### Preparation of *E. coli* samples for super resolution microscopy

For fluorescence microscopy analysis, single colonies were used to inoculate LB medium and grown overnight at 37°C with shaking. Cultures carrying plasmids were supplemented with the appropriate antibiotics. Overnight cultures were diluted 1:100 into fresh LB medium and grown for approximately 2 h at 37°C until exponential phase. Cultures were subsequently diluted to an OD_600_of 0.02 and grown at the indicated temperatures (37°C or 25°C) until mid-log phase (OD_600_ 0.4–0.6). For chemical perturbation experiments, exponentially growing cultures were treated with the indicated compounds under the following conditions unless otherwise stated: 5% (v/v) 1,6-hexanediol for 5 min, 50 μg mL⁻¹ rifampicin for 15 min, or 1 mM IPTG for 60–90 min for the induction of mCherry-ParB. For fixation, 2 mL of cell suspension was harvested by centrifugation at 5,000 ×g for 2 min and resuspended in 4% methanol-free paraformaldehyde (PFA) prepared in 1× PBS. Cells were fixed for 15 min at room temperature and subsequently washed once with 1× PBS. DNA was stained by incubation with Hoechst 33342 (Thermo Fisher Scientific) at a final concentration of 20 μg mL⁻¹ in 1× PBS for 1 h. Halo-tagged strains were additionally labelled with 100 nM Janelia Fluor 549 HaloTag Ligand (Promega) for 1 h. Excess dye was removed by washing cells three times with 1× PBS. Following washing, cells were resuspended in 100 μL PBS. A 20 μL aliquot was centrifuged and resuspended in 10 μL Vectashield mounting medium (Vector Laboratories). Five microliters of the sample were spotted onto 1.5% agarose pads and covered with 1.5 thickness coverslips prior to imaging. All microscopy samples were prepared using biological triplicates.

### 3D Structured Illumination Microscopy (3D-SIM)

3D-SIM imaging was performed as follows. Fixed *E. coli* cells prepared for super resolution microscopy were imaged using a Deltavision OMX-BLAZE microscope system (Applied Precision) equipped with a 60×/NA 1.42 Plan Apo N oil immersion objective, four liquid-cooled sCMOS cameras, and laser lines at 405, 488, 568, and 642 nm. Images were acquired as z-stacks using a 512 × 512 pixel field of view with 1 μm total z-range and 0.125 μm z-step size, 9 sections. Each optical section was acquired using a striped illumination pattern rotated by −60°, 0°, and +60° and shifted through five phase positions. Hoechst 33342-stained DNA was imaged using the 405 nm laser (15% power, 40 ms exposure). mEGFP-labelled proteins were imaged using the 488 nm laser at either 5% or 20% power with 20 ms or 40 ms exposure times, respectively. Halo-tagged proteins and mCherry-ParB were imaged using the 568 nm laser at 20% power with a 60 ms exposure time. For all acquisitions the Cargille LASER Liquid oil with n=1.5160 +/- 0.0002, 589.3nm (Code 5610) was used.

Raw images were reconstructed and channels were aligned using the softWoRx software package (Applied Precision). Maximum intensity z-projections were generated using five of the nine z-planes and image panels were assembled using OMERO^8^. Overview images were displayed at 2× magnification and zoomed-in regions at 5× magnification. Where both overview and zoomed-in images are shown, the area corresponding to the magnified view is indicated by a white box in the overview image. For each of three independent biological replicates, three to five fields of view were acquired per sample, resulting in at least hundred cells analysed per replicate. Laser power and exposure times were kept constant between biological replicates and different experiments if not indicated differently.

### Quantification of super resolution microscopy images

Maximum-intensity z-projections generated from reconstructed 3D-SIM images as described above were quantified using a custom Python pipeline (Python v3.13) based on NumPy^9^, SciPy^10^ and scikit-image^11^. The reconstructed pixel size of 0.0397 μm was used to convert pixel-based measurements into physical units. Unless otherwise stated, Hoechst fluorescence was assigned to channel 0 and mEGFP fluorescence to channel 1, with mCherry-P1-ParB, mKOk or Janelia Fluor 549 fluorescence assigned to channel 2 for three-colour datasets.

Because the number of detected CsdA foci was sensitive to acquisition intensity, all CsdA quantifications were performed exclusively using images acquired with the shorter mEGFP exposure (488 nm laser, 5% power, 20 ms exposure). DbpA, RhlB, RhlE and SrmB, which were imaged using the higher exposure setting (488 nm laser, 20% power, 40 ms exposure), were quantified separately.

Cells were segmented using Cellpose v4.2.1.1 with the Cellpose-SAM model and GPU acceleration^12,13^. Segmentation was generally performed using the mEGFP channel, which delineated the cytoplasmic cell volume. For endogenously Halo-tagged colocalisation strains, the channel providing the most complete cell outline was used for segmentation: the CsdA-HaloTag/Janelia Fluor 549 channel when the co-expressed mEGFP-tagged marker showed a punctate distribution, or the mEGFP channel when the marker showed a cytoplasmic distribution. Segmentation masks contacting the image boundary were excluded, and masks with a cross-sectional area outside 300–60,000 pixels (0.47–94.6 μm²); the lower bound removes sub-cellular debris, whereas the upper bound excludes only rare merged-cell segmentation artefacts, as all genuine cells fell well below it (maximum 10.95 µm²).

Fluorescent foci were detected in the mEGFP channel using Spotiflow v0.6.5^14^. The base Spotiflow model was fine-tuned using manually annotated CsdA images to generate a detector specific for CsdA foci. During training annotation, greater confidence was assigned to foci occurring as spatially clustered signals within cells. Separate detectors for SrmB and RhlB, which also formed discrete foci, were fine-tuned from the same base model. Detection was performed on raw, non-renormalised fluorescence intensities because intensity renormalisation resulted in systematic under-detection of foci in cells with high fluorescence intensity. Candidate foci were initially predicted using a probability threshold of 0.02 and subsequently filtered to retain detections with a probability ≥0.20. Detected foci were assigned to individual cells using the corresponding segmentation masks. DbpA and RhlE showed diffuse cellular distributions without bona fide foci. Because application of the CsdA-trained detector to these strains primarily detected shot-noise speckles in cells with low fluorescence intensity, these detections were considered imaging artefacts and were not interpreted as foci or included in quantitative focus comparisons.

For each cell, the number of detected foci, median focus area and mean focus intensity were determined. The percentage of cells containing foci was calculated as the fraction of valid segmented cells exceeding the indicated focus-number threshold, and the mean number of foci per cell was calculated across all valid cells.

For visualisation and spatial analysis of the *parS*-labelled *rrnD* locus, mCherry-P1-ParB foci were detected in the red channel using a local-maximum detection approach. Candidate maxima were required to exceed the local background by more than eight median absolute deviations and to be separated by at least four pixels. Under these conditions, ParB foci were detected in ≥98% of cells carrying the *parS*-labelled locus, whereas <6% of cells in the parental strain lacking a *parS* insertion scored positive. CsdA–*rrnD* proximity was quantified as the distance between each CsdA focus and the nearest ParB focus. Observed distances were compared with a within-cell random distribution generated from five random-position draws per CsdA focus. Radial enrichment profiles were calculated from the same distance measurements as the ratio between observed and random densities in consecutive 50 nm shells. Confidence intervals were determined by 500 bootstrap resamplings over cells. For radial-density representations, mean fluorescence intensity in the indicated channel was calculated in concentric 50 nm shells centred on the ParB focus and normalised to the mean fluorescence intensity of the corresponding cell, such that a value of one represents the cell-average fluorescence intensity.

Colocalisation of CsdA with the ribosome biogenesis and transcription-associated proteins RpoC, NusB, ObgE, RplA and RpsB was analysed separately for plasmid-expressed mKOk-tagged proteins and endogenously tagged mEGFP/HaloTag strains because the two datasets differed in tag chemistry, channel configuration and experimental controls. Channel identities were verified from the corresponding image datasets prior to analysis. Colocalisation was primarily quantified using a matched-null enrichment coefficient. For each CsdA focus, marker-channel fluorescence within the CsdA focus footprint was divided by the mean marker fluorescence measured at random positions within the same cell matched for distance to the nucleoid. This matching controlled for enrichment arising solely from the shared tendency of CsdA and the analysed proteins to localise to the nucleoid and therefore measured marker enrichment at CsdA foci above that expected from co-occupancy of the same cellular compartment. The CsdA strain lacking the corresponding fluorescent marker was used to define the background enrichment level.

Quantified percentages of cells containing foci and numbers of foci per cell were plotted in GraphPad Prism as mean ± SD. Statistical significance was assessed using ordinary one-way ANOVA followed by Tukey’s multiple-comparisons test or Fisher’s LSD test, as indicated. Significance for selected pairwise comparisons is indicated as follows: ns, *P* ≥ 0.1234; *, *P* < 0.0332; **, *P* < 0.0021; ***, *P* < 0.0002; ****, *P* < 0.0001.

### Growth curves of endogenous mutant strains

For all tested untagged or mEGFP-tagged CsdA mutant strains, three individual colonies were used to set up overnight cultures in LB medium and incubated at 37°C with shaking. Overnight cultures were diluted 1:100 into fresh LB medium and grown for approximately 1.5 h at 37°C with shaking. Cultures were adjusted to an OD_600_ of 0.01 in LB medium and used to inoculate wells of a 96-well flat-bottom plate containing LB medium at a starting OD600 of 0.001 in a final volume of 150 μL per well. Plates were incubated at 25°C or 37°C in a SPECTROstar Nano plate reader (BMG Labtech) with double-orbital shaking (500 rpm) and OD_600_measurements every 30 min for 48 h. In parallel, cultures diluted to an OD_600_ of 0.02 in 25 mL LB medium were incubated at 25°C with shaking and sampled for western blot analysis and fluorescence microscopy at an OD_600_ of 0.4-0.6.

### Complementation assay of Δ*csdA* strain

To assess the ability of plasmid-expressed CsdA variants to rescue the cold-sensitive growth defect of a Δ*csdA* strain, previously also referred to as Δ*deaD* strain, *E. coli* K-12 MG1655 Δ*csdA* carrying plasmids encoding CsdA^WT^-mEGFP, CsdA^ΔIDR1/2^-mEGFP, or mEGFP alone under constitutive promoters of varying strength were compared to chromosomally tagged CsdA-mEGFP and Δ*csdA* control strains. For each strain, three independent colonies were used to inoculate 5 mL LB medium supplemented with kanamycin when required. Overnight cultures were grown at 37°C with shaking. The following day, cultures were diluted 1:100 into fresh LB medium and incubated for 1.5 h at 37°C with shaking to obtain exponentially growing cells. For growth measurements, cultures were diluted to an OD_600_ of 0.01 and used to inoculate 96-well plates containing LB medium (with or without kanamycin as appropriate) at a final starting OD_600_ of 0.001. Wells contained 150 μL culture volume and were prepared in biological triplicates together with medium-only controls for blank subtraction. Plates were incubated at 25°C in a microplate reader with double-orbital shaking (500 rpm), and OD600 measurements were recorded every 30 min for 48 h. Growth curves were generated from blank-subtracted absorbance values and used to compare the growth of the different complementation strains. In parallel, larger-volume assay cultures were prepared for protein expression analysis. Exponentially growing cultures were diluted to an OD_600_ of 0.02 in 25 mL LB medium and incubated at 25°C with shaking. Aliquots were collected at an OD_600_ of 0.4- 0.6 for subsequent western blot analysis to correlate growth rescue with expression levels of the CsdA-mEGFP variants.

For both growth curves and rescue assay absorbance values were blank-subtracted and resulting growth curves were plotted in Prism (GraphPad). Growth curves are shown as the mean ± SD of three independent biological replicates. The time required to reach half of the maximal OD_600_ (T_50_) was determined for each biological replicate using a custom R script. Briefly, negative OD_600_ values resulting from blank subtraction were set to zero, and only measurements acquired up to the first occurrence of the maximal OD_600_ were included in the analysis to exclude the decline in stationary phase. T_50_ was calculated by linear interpolation between the two consecutive measurements bracketing 50% of the maximal OD_600_. T_50_ values are presented as the mean ± SD of three independent biological replicates and plotted using Prism (GraphPad).

Statistical significance was assessed using an ordinary one-way ANOVA with Tukey’s multiple-comparisons test. Significance for selected pairwise comparisons are indicated as follows: ns, P ≥ 0.1234; P < 0.0332(*); P < 0.0021(**); P < 0.0002(***); P < 0.0001(****).

### Fluorescents immunoblot analysis

To compare expression levels of chromosomally encoded CsdA-mEGFP variants and proteins expressed from plasmids with varying expression strength, cells were grown to mid-log phase (OD_600_ 0.4–0.6) and harvested by centrifugation of 2 mL culture at 5,000 ×g for 2 min at room temperature. Cell pellets were snap-frozen in liquid nitrogen and stored at −20°C until further processing.

For lysis, pellets were thawed on ice and resuspended in lysis buffer (1x phosphate-buffered saline (PBS), 0.2 mg mL^−1^ lysozyme, 20 µg mL^−1^ DNase I, 1 mM PMSF, 1 mM MgCl_2_). The volume was adjusted according to cell density (100 μL for a culture with an OD_600_ of 0.5). Samples were incubated for 15 min at 30°C to allow enzymatic lysis. Lysate was mixed with an equal volume of 2x LI-COR Orange Protein Sample Buffer supplemented with DTT, sonicated for 5 min in a water bath sonicator, and heated at 95°C for 7 min. Subsequently, 6 μL of each sample were loaded onto 4–12% Bis-Tris gradient gels (NuPAGE™, Thermo Fisher Scientific) together with a prestained molecular weight marker (PageRuler™ Plus Prestained Protein Ladder, Thermo Fisher Scientific).

Electrophoresis was performed in MOPS running buffer at 180 V for 50–60 min. Proteins were transferred onto Immobilon®-FL PVDF membranes (Millipore) using a wet transfer system at 50 V for 60 min in using Tris-glycine buffer (25 mM Tris pH 8.3, 192 mM glycine). Following transfer, membranes were dried overnight between filter papers.

Total protein levels were assessed prior to immunodetection using Revert™ 520 Total Protein Stain (LI-COR). Membranes were rehydrated in methanol, equilibrated in TBS, stained for 5 min at room temperature, washed according to the manufacturer’s instructions, and imaged immediately at 520 nm using an ODYSSEY® Imaging System. Following imaging, membranes were destained and prepared for antibody detection. Membranes were blocked for 1 h at room temperature in Intercept® TBS Blocking Buffer (LI-COR). Tagged proteins were detected using a rabbit anti-GFP primary antibody (Thermo Fisher Scientific A11122; 1:3’000 dilution) or mouse anti-Halo primary antibody (Chromotek, 28A8; 1:5’000 dilution) in blocking buffer supplemented with 0.2% Tween-20. Membranes were incubated with the primary antibody for 1.5 h at room temperature, washed four times for 5 min with TBS-T (20 mM Tris-HCl pH 7.5, 150 mM NaCl, 0.2% (w/v) Tween20)), and subsequently incubated for 1 h at room temperature with IRDye® 800CW donkey anti-rabbit secondary antibody or IRDye® 680RD donkey anti-mouse (LI-COR; 1:20,000 dilution) diluted in blocking buffer containing 0.2% Tween-20 and 0.02% SDS. All secondary antibody incubations and subsequent washes were performed protected from light. After four additional washes with TBS-T and a final rinse with TBS, membranes were imaged using an ODYSSEY® Imaging System. Western blots were performed in triplicates.

Revert™ 520 total protein staining was used as a loading control to assess comparable sample loading. Protein-specific band fluorescence intensities were quantified using Empiria Studio® software (LI-COR). Three independent western blots were analysed for each experiment. Depending on the experiment, protein abundance was either reported directly as fluorescence intensity in arbitrary units (AU) or expressed as relative protein abundance after normalisation to the CsdA^WT^-mEGFP reference sample grown at the indicated temperature (25°C or 37°C). Quantified values from the three independent experiments were plotted as mean ± SD.

For the *ΔcsdA* growth-rescue experiment, CsdA^WT^-mEGFP and CsdA^ΔIDR1/2^-mEGFP protein levels obtained from the graded-expression series were normalised to endogenous CsdA^WT^-mEGFP abundance at 25°C. Mean relative protein abundance across the three independent western blots was then related to the corresponding T_50_ values derived from the growth-curve analysis and displayed as an XY scatter plot, with relative protein abundance plotted on a logarithmic scale. Western blots of strains carrying both Halo- and mEGFP-tagged proteins were used solely to verify expression and the presence of both tagged proteins and were therefore not subjected to quantitative analysis.

If applicable statistical significance was assessed using an ordinary one-way ANOVA with Fisher’s LSD test and 95% confidential interval. Significance for selected pairwise comparisons are indicated as follows: ns, P ≥ 0.1234; P < 0.0332(*); P < 0.0021(**); P < 0.0002(***); P < 0.0001(****).

### Whole cell mass spectroscopy (MS)

For quantitative proteomic analysis, *E. coli* strains were grown in biological triplicates. Overnight cultures were incubated in LB medium at 37°C with shaking. Cultures were diluted 1:100 into fresh LB medium and grown for 2 h at 37°C with shaking. Pre-cultures were used to inoculate assay cultures at a starting OD_600_ of 0.02, which were subsequently incubated at the indicated temperatures (37°C, 25°C and/or 20°C). Cells were harvested during mid-log phase (OD_600_ 0.4–0.6) by collecting approximately 1 × 10⁹ cells per sample (based on OD_600_ conversion where OD_600_ = 1 corresponds to 8 × 10⁸ cells mL⁻¹) by centrifugation at 4,500 ×g for 5 min at 4°C. Cell pellets were washed twice with ice-cold PBS, snap-frozen in liquid nitrogen, and stored at −75°C until further processing.

For protein extraction, cell pellets were resuspended in 80 μL freshly prepared lysis buffer containing 5% (w/v) sodium dodecyl sulfate (SDS), 10 mM tris(2-carboxyethyl)phosphine (TCEP), and 100 mM triethylammonium bicarbonate (TEAB). Cells were lysed using a PIXUL Multi-Sample Sonicator (Active Motif) for 20 min at 15°C (pulse 50 N, PRF 1 kHz, burst rate 20 Hz), followed by incubation at 95°C for 10 min. Protein concentration was determined using a tryptophan-based fluorescence assay (Infinite M Plex, Tecan). Protein lysates were alkylated by addition of iodoacetamide (IAA) to a final concentration of 20 mM and incubation for 30 min at room temperature protected from light. A total of 20–50 μg of alkylated protein lysate was digested with trypsin and purified using S-trap columns. Peptide concentration was determined using an UV-based assay (Infinite M Nano, Tecan), and samples were adjusted to a final peptide concentration of 250 ng μL⁻¹ prior to mass spectrometry analysis. Samples were analysed in biological triplicates.

### Affinity Purification Mass Spectrometry (AP-MS)

For affinity purification mass spectrometry, *E. coli* K-12 MG1655 strains carrying endogenously mEGFP-tagged DEAD-box proteins were compared to a wild-type control strain expressing mEGFP from plasmid pMH2581 (pEB1(J23108)-mEGFP-A206K). Cultures were prepared as described for relative quantitative mass spectrometry analysis with minor adaptations. In brief, overnight cultures were diluted into fresh LB medium (supplemented with kanamycin for the mEGFP control strain) to a starting OD_600_ of 0.02 and grown in biological triplicates at the indicated temperatures (37°C or 25°C). Cells were harvested at OD_600_ 0.4– 0.6 by collecting approximately 2.5 × 10⁹ cells per sample by centrifugation at 4,500 ×g for 5 min at 4°C. Cell pellets were washed twice with ice-cold PBS, snap-frozen in liquid nitrogen, and stored at −75°C until further processing.

For GFP-based affinity purification, cell pellets were resuspended in 100 μL freshly prepared lysis buffer containing 50 mM Tris-HCl pH 7.5, 150 mM NaCl, 2 mM MgCl₂, 0.5% (w/v) NP-40, 1 mM PMSF, and protease inhibitors. Cells were lysed using a PIXUL Multi-Sample Sonicator (Active Motif) for 20 min at 15°C (pulse 50 N, PRF 1 kHz, burst rate 20 Hz). Lysates were diluted with 400 μL ice-cold dilution buffer containing 50 mM Tris-HCl pH 7.5, 150 mM NaCl, 2 mM MgCl₂, 1 mM PMSF, and protease inhibitors. Diluted lysates were centrifuged at 5,000 ×g for 2 min and clarified supernatant was incubated with 25 μL equilibrated GFP-Trap magnetic agarose beads (Chromotek) for 2 h at 4°C with rotation. Beads were separated from the lysate using a magnetic rack, and unbound fractions were removed. Beads were washed four times with ice-cold wash buffer (50 mM Tris-HCl pH 7.5, 150 mM NaCl, 2 mM MgCl₂, 0.05% (w/v) NP-40, 1 mM PMSF, and protease inhibitors), followed by two additional washes with dilution buffer without NP-40. Bound proteins were eluted from the beads in 45 μL elution buffer containing 5% (w/v) SDS, 1 mM TCEP, and 100 mM TEAB by incubation at 95°C for 10 min.

Eluted proteins were alkylated by addition of IAA to a final concentration of 20 mM and incubation for 30 min at room temperature protected from light. Proteins from the complete eluate were purified and digested with the SP3 approach using a Fluent liquid handling platform (Tecan Group Ltd)^15^. In brief, Speed BeadsTM (GE Healthcare) were mixed 1:1, rinsed with water and diluted to the 8 μg µL^−1^ stock solution. Samples were adjusted to the final volume of 45 µL and 10 µL of the beads stock solution was added to them. Proteins were bound to the beads by addition of 125 µL of 100% acetonitrile to the samples, which were then incubated for 8 min at RT with a gentle agitation (200 rpm). After, samples were placed on a magnetic rack and incubated for 5 minutes. Supernatants were removed and discarded. The beads were washed twice with 160 µL of 70% (v/v) ethanol and once with 160 µL of 100% acetonitrile. Samples were placed off the magnetic rack and 50 µL of digestion mix (10 ng µL^−1^ of trypsin in 50 mM triethylammonium bicarbonate) was added. Tryptic digestion proceeded for 12 h at 37°C. After digestion samples were placed back on the magnetic and incubated for 5 minutes. Supernatants containing peptides were collected and dried under vacuum. AP-MS experiments were performed in biological triplicates.

### SWATH-MS data acquisition for MS and AP/MS

Peptides were re-suspended in 0.1% aqueous formic acid and 0.1 µg of peptides subjected to LC–MS/MS analysis using an Orbitrap Exploris 480 MassSpectrometer fitted with an EASY-nLC 1200 (both Thermo Fisher Scientific) and a custom-made column heater set to 60°C. Peptides were resolved using a RP-HPLC column (75 μm × 30 cm) packed in-house with C18 resin (ReproSil-Pur Saphire, 1.5μm resin; Dr. Maisch GmbH) at a flow rate of 0.2 μL min^−1^. Separation of peptides was achieved using the following gradient from Buffer A (0.1% formic acid in water) to Buffer B (80% acetonitrile, 0.1% formic acid in water): starting from 2% to 8% Buffer B in 5 min, to 25% Buffer B in 25 min, to 35% Buffer B in 10 min. The mass spectrometer was operated in DIA mode with a cycle time of 3 seconds. MS1 scans were acquired in the Orbitrap in centroid mode at a resolution of 120,000 FWHM (at 200 m/z), a scan range from 350 to 910 m/z, normalised AGC target set to 300 % and maximum ion injection time mode set to 50 ms. MS2 scans were acquired in the Orbitrap in centroid mode at a resolution of 15,000 FWHM (at 200 m/z), precursor mass range of 400 to 900, quadrupole isolation window of 12 m/z with 1 m/z window overlap, a defined first mass of 120 m/z, normalised AGC target set to 3000 % and a maximum injection time of 22 ms. Peptides were fragmented by HCD (Higher-energy collisional dissociation) with collision energy set to 28 % and one microscan was acquired for each spectrum.

### SWATH-MS data analysis for MS and AP/MS

The acquired raw files were searched using SpectroNaut default settings (Biognosys) against a *E. coli* K12 MG1665 database (consisting of 4,390 protein sequences downloaded from www.uniprot.org on 2022/02/22) and 393 commonly observed contaminants using the following search criteria: full tryptic specificity was required (cleavage after lysine or arginine residues, unless followed by proline); 3 missed cleavages were allowed; carbamidomethylation (C) was set as fixed modification; oxidation (M), N-acetlyation (N-term) were applied as variable modifications. The raw quantitative data was further statically analysed using ProteoFlux, an open-source and reproducible workflow for quantitative proteomics data analysis (https://doi.org/10.5281/zenodo.18640999). Prior to quantification, precursor intensities were filtered based on run-level posterior error probability (PEP) and q-value thresholds (≤ 0.01), and very low-intensity signals (below a threshold of 2) were treated as left-censored measurements. Precursor-level intensities exported from Spectronaut were aggregated to peptide-level quantities and subsequently summarized to protein abundances using a simple summarisation-based approach, followed by median-based sample normalisation and statistical analysis using linear models with empirical Bayes moderation. AP-MS data were further normalised to GFP prior data visualisation by R (R Studio). For AP-MS and whole-cell MS -log_10_(q-value) was plotted against the log_2_(fold change) of proteins that were detected in all 3 biological replicates, showing at least 3 consistent peptides.

For the whole cell MS, the estimated cellular protein concentration was calculated based on the measured fold change between 37°C and 25°C samples, the previously reported protein copy numbers per cell at 37°C and *E. coli* cell volume measurements of 4 fL^16,17^.

### Protein expression and purification

Recombinant proteins were expressed in *E. coli* Lemo21(DE3) transformed with pETMCN-based expression plasmids under the respective antibiotic selection. Expression cultures were launched by diluting overnight cultures grown in LB containing 1 % D(+)-glucose to a starting OD_600_ of 0.05 in Terrific-Broth (TB) medium supplemented with the appropriate antibiotics. After approx. 3 h at 37°C, protein expression was induced at an OD_600_ of 0.6-0.8 by the addition of 0.2 mM IPTG. Cells were further grown at 20°C overnight and were harvested by centrifugation at 5,000 ×g for 15 min at 4°C. Pellets were snap frozen in liquid nitrogen and stored at -75°C until further processing.

Pellets were resuspended in 30 mL lysis buffer (25 mM Tris-HCl pH 7.5 or pH 8.0, 1 M NaCl, 5 % glycerol (w/v), 20 mM imidazole, protease inhibitors, DNase I, and RNase A) per 1 L expression culture. Cells were either lysed by sonication on ice or by pressure homogenisation using EmulsiFlex (Avestin). Bacterial lysates were cleared by centrifugation at 20,000 ×g for 30 min at 4°C and filtration through a 0.45 µm filter. His_10_-tagged proteins were purified by immobilized metal affinity chromatography (IMAC) using Ni-NTA Sepharose gravity flow column and elution with high concentrations of imidazole. Overnight affinity purification tag was removed by 3C Pressision protease cleavage and simultaneous dialysis into 25 mM Tris-HCl pH 7.5 or pH 8.0, 1 M NaCl, 5 % glycerol (w/v), 2 mM MgCl_2_ and 0.5 mM 2-ME. Uncleaved proteins and cleaved affinity tags were removed by reverse IMAC. Proteins were further purified by size exclusion chromatography (SEC) on a HiLoad^®^ 16/600 Superdex^®^ 200 pg column (Cytiva) using an ÄKTA pure (GE Life Sciences) into the protein storage buffer (25 mM Tris-HCl pH 7.5 or pH 8.0, 1 M NaCl, 10 % glycerol (w/v), 2 mM MgCl_2_ and 0.5 mM 2-ME). During the protein purification main fractions of the IMAC, reverse-IMAC, SEC and the final protein pool were analysed by Sodium Dodecyl Sulphate Polyacrylamide Gel Electrophoresis (SDS-PAGE) under reduced conditions followed by a Coomassie blue staining. The AcuteBand Prestained Protein Ladder (Lubio Science) was loaded as a molecular weight reference. Final protein pools were up concentrated using Amicon^®^ Ultra Centrifugal Filter (Merck) with an appropriate molecular weight cut off (MWCO). Protein concentration was calculated based on the molecular weight (MW), specific molar extinction coefficient (ε280) and the absorbance measured at 260 nm, 280 nm and 333 nm by NanoDrop (Thermo Scientific). Concentrated proteins were aliquoted, snap frozen in liquid nitrogen and stored at -75°C until further usage. For all purified proteins, protein integrity was assessed by analysing 0.5 µg total protein per lane on an SDS-PAGE.

An overview of the protein characteristics, including the MW, ε280, isoelectric point (PI), net charge, and storage buffer information are listed in (Extended Data Table 2).

### Chemical Labelling of Purified Proteins

Purified untagged proteins were chemically labelled at the N-terminus as described previously with minor adaptations^18^. Briefly, proteins were adjusted to a concentration of 50 μM in protein storage buffer containing 25 mM Tris-HCl pH 7.5, 1 M NaCl, 2 mM MgCl₂, 10% (w/v) glycerol, and 0.5 mM β-mercaptoethanol. Proteins were exchanged into labelling buffer consisting of 25 mM KH₂PO₄/K₂HPO₄ pH 6.5, 1 M NaCl, 2 mM MgCl₂, 10% (w/v) glycerol, and 0.5 mM β-mercaptoethanol using Zeba Spin desalting columns (Thermo Fisher Scientific; 7 kDa MWCO). N-terminal labelling was performed by adding a four-fold molar excess of ATTO488 or ATTO565 NHS-ester dye (ATTO-TEC) dissolved in DMSO. Labelling reactions were incubated for 1 h at room temperature protected from light. Excess unbound dye was removed by buffer exchange using Zeba Spin desalting columns equilibrated with protein storage buffer. Remaining free dye was further removed by repeated concentration and dilution steps using Amicon Ultra centrifugal filters (Merck; 30 kDa MWCO).

Protein concentration and degree of labelling were determined by UV/Vis spectrophotometry using a NanoDrop instrument (Thermo Fisher Scientific) by measuring absorbance at 280 nm and 500 nm. Calculations were performed considering the protein-specific molar extinction coefficient, ATTO dye extinction coefficient, correction factor at 280 nm, and molecular weight of the labelled protein. For fluorescence-based in vitro condensation assays, fluorescently labelled protein was mixed with unlabelled protein at a final fraction of approximately 2% labelled protein for visualisation.

### In vitro condensation assays

In vitro condensation assays were performed as described previously with minor adaptations ^1^. Purified untagged proteins were diluted to ten times the highest assay concentration in the corresponding protein storage buffer containing 25 mM Tris–HCl (pH 7.5 or 8.0), 1 M NaCl, 10% (w/v) glycerol, 2 mM MgCl₂ and 0.5 mM β-mercaptoethanol. Proteins were supplemented with 2% fluorescently labelled protein for visualisation.

To determine the in vitro saturation concentration (*c*_sat_), proteins (including all bacterial DDXs and CsdA variants) were serially diluted in a two-fold dilution series using the corresponding protein storage buffer. For condensation assays, 2 μL of protein solution was mixed with 18 μL of assay master mix in black, optically clear, flat-bottom, ultra-low-attachment-coated 384-well PhenoPlates (Revvity). Unless indicated otherwise, the final assay conditions were 30 mM HEPES–NaOH (pH 7.5), 150 mM NaCl, 2 mM MgCl₂, 1% (w/v) glycerol, 0.5 mg mL⁻¹ BSA and either 0.05 mg mL⁻¹ polyuridylic acid (poly(U)), 0.05 mg mL⁻¹ total *E. coli* RNA or no RNA. Final protein concentrations ranged from 8.0 to 0.25 μM. Plates were centrifuged briefly for 20 s at 10 × *g* and incubated at 25°C for 30 min prior single time-point image acquisition.

For time- and temperature-dependent condensation measurements, reactions were assembled as described above, with solutions pre-warmed to the indicated temperature. Reactions were incubated at 25°C, 30°C or 37°C and imaged every 5 min for 1 h.

For time-resolved assays examining the effects of RNA and ATP regeneration, untagged wild-type CsdA, CsdA^DQAD^ or CsdA^ΔIDR1/2^ was used at a final concentration of 2 μM and supplemented with 2% of the corresponding ATTO488-labelled protein. Reactions contained 30 mM HEPES–NaOH (pH 7.5), 150 mM NaCl and either 0.05 mg mL⁻¹ poly(U), 0.05 mg mL⁻¹ total *E. coli* RNA or no RNA. All reactions additionally contained a creatine kinase-based ATP-regeneration mixture (CMK) at a final concentration of 2 mM ATP, 2 mM MgCl₂, 10 mM creatine phosphate and 3.5 U mL⁻¹ creatine kinase. ATP-negative controls contained an otherwise equivalent CMK mixture prepared without ATP. Reactions were incubated at 25°C and imaged every 10 min for 1 h.

For co-condensation assays, wild-type CsdA and the indicated DEAD-box proteins (SrmB, DbpA, RhlE and RhlB) or ribosome biogenesis factors (ObgE and HrpA) were fluorescently labelled with ATTO488 and ATTO565, respectively. Proteins were assayed at a final concentration of 2 μM either individually or in mixtures containing 2 μM CsdA and 2 μM of the indicated partner protein. Reactions were assembled under the standard condensation conditions described above and contained 0.05 mg mL⁻¹ poly(U). Samples were incubated at 25°C for 30 min prior to imaging.

Images were acquired automatically using the Nikon JOBS system integrated into Nikon NIS-Elements software on a Nikon Ti2-E inverted spinning-disk confocal microscope equipped with a CrestOptics X-Light V3 spinning-disk module, a seven-channel Lumencor CELESTA light source, a 40×/0.95 NA Plan Apo Lambda air objective and a 10.2-megapixel back-illuminated sCMOS camera. Samples were imaged in spinning-disk mode using the green fluorescence channel and, for co-condensation assays, the corresponding red fluorescence channel. Four fields of view were acquired per condition in each of three independent experiments. Laser power and exposure times were kept constant across conditions and independent experiments. Image processing and visualisation were performed using the Open Microscopy Environment Remote Objects platform (OMERO)^8^. Representative images show selected conditions from one of the three independent experiments.

Fluorescence images were analysed using CellProfiler^19^. For condensation analysis, the mean fluorescence intensity and fluorescence intensity standard deviation were determined for each image. A condensation index, corresponding to the coefficient of variation (CV), was calculated in R by dividing the fluorescence intensity standard deviation by the mean fluorescence intensity of each image. Higher values indicate greater spatial heterogeneity of the fluorescence signal and increased condensation, whereas lower values indicate a more homogeneous fluorescence distribution. For concentration-dependent assays, the mean condensation index ± SD was plotted against protein concentration to estimate the *c*_sat_ of each DEAD-box protein or CsdA variant. Temperature-dependent condensation dynamics were assessed by plotting the CV over time at each temperature. For the CMK assays, condensation dynamics were similarly plotted over time to compare CsdA variants in the presence or absence of ATP and in reactions containing poly(U), total *E. coli* RNA or no RNA.

For co-condensation assays, CV were calculated independently for the CsdA and partner-protein fluorescence channels to assess changes in spatial fluorescence heterogeneity upon mixing. Colocalisation between CsdA and the respective partner protein was quantified using Pearson’s correlation coefficient. CsdA condensates were identified from the ATTO488 channel, and Pearson correlation coefficients between the ATTO488 and ATTO565 fluorescence intensities were calculated within the segmented condensate regions for each image. Values from all identified objects were averaged to obtain one value per condition for each independent experiment. Data are presented as the mean ± SD from three independent experiments (N = 3).

Condensate morphology was quantified from CellProfiler-segmented objects using the circularity measurement, FormFactor, calculated as ((4π*πr^2^)/(2πr)^2^), where values approaching 1 indicate increasingly circular objects and lower values indicate deviation from a circular shape. To minimize bias from pixelation of small objects, condensates with an area <100 pixels² were excluded from the analysis. FormFactor analysis was performed after 40 min of incubation under the indicated RNA and ATP conditions. For each field of view, the median FormFactor of all detected condensates was calculated. Median values from four fields of view were averaged to obtain one value per independent experiment, and data are presented as the mean ± SD from three independent experiments (N = 3).

Statistical significance was assessed using an ordinary one-way ANOVA with Tukey’s multiple-comparisons test. Significance for selected pairwise comparisons are indicated as follows: ns, P ≥ 0.1234; P < 0.0332(*); P < 0.0021(**); P < 0.0002(***); P < 0.0001(****).

### ATPase assays

Enzymatic activity of the wild-type CsdA and truncation variants were determined in an indirect photometric ATPase assay as described before^20,21^. In short, two 2-fold concentrated master mixes were prepared allowing controlled reaction initiation and kinetic absorbance measurements. First, to assemble master mix A prediluted proteins in protein storage buffer were mixed with or without poly(U) as RNA substrate, BSA and glycerol in tenfold concentrated ATPase assay buffer with NaCl (300 mM HEPES-NaOH pH 7.5, 1 M NaCl, 20 mM MgCl_2_). Secondly, adenosine 5′-triphosphate (ATP) pH 7.5, dithiothreitol (DTT), phosphoenolpyruvicacid (PEP) pH 7.5, reduced nicotinamide adenine dinucleotide (NADH) and pyruvate kinase and lactic dehydrogenase enzymes (PKL) were mixed in tenfold concentrated ATPase assay buffer without NaCl (300 mM HEPES-NaOH pH 7.5, 20 mM MgCl_2_) for master mix B. ATP hydrolysis reactions were launched by mixing equal volumes of master mix A and B in a 384-well black, clear flat bottom plate (Corning) resulting in the final assay condition of 1 µM DDX, none or 0.5 mg mL^−1^ poly(U) in 30 mM HEPES-NaOH pH 7.5, 100 mM NaCl, 2 mM MgCl_2_, 10 % glycerol (w/v), 0.33 mg mL^−1^ BSA, 2.5 mM ATP, 10 mM DTT, 6 mM PEP, 1.2 mM NADH and 11-25 units mL^−1^ PKL. Reactions with and without RNA substrate were set up. NADH absorption was monitored with a Spark Multi-Mode plate reader (Tecan) at 340 nm in approx. 70 s intervals for 34 cycles.

Pathlength corrected absorbance values in OD cm^−1^ were used to calculate the NADH concentration using the Beer-Lambert law using ε_340_ of 6220 M^−1^ cm^−1^, and a pathlength of 1 cm. To determine the initial reaction kinetics NADH concentration of the triplicates was plotted against the assay time using Prism (GraphPad). A linear regression fit was used to determine the initial slope of the reaction representing the NADH consumption or ATP consumption interchangeably. ATP consumption rate per protein was determined considering the final protein concentration. Experiments were performed in technical triplicates and repeated in three independent experiments. Mean ATP consumption rate ± SD of three independent experiments (N=3).

Statistical significance was assessed using an ordinary one-way ANOVA with Tukey’s multiple-comparisons test. Significance for selected pairwise comparisons are indicated as follows: ns, P ≥ 0.1234; P < 0.0332(*); P < 0.0021(**); P < 0.0002(***); P < 0.0001(****).

### Polysome profiling

Polysome profiles of *E. coli* strains grown at different temperatures were analysed as described previously with minor adaptations^22^. Briefly, 10–40% or 10–25% linear sucrose gradients were prepared using a Gradient Master 108 (Biocomp) with equal volumes of high-and low-sucrose buffer containing 10 mM Tris-HCl pH 7.5, 60 mM KCl, 50 mM NH₄Cl, 10 mM MgCl₂, 1 mM DTT, and either 10%, 25%, or 40% (w/v) sucrose. Gradients were prepared and stored overnight at 4°C prior to use.

For polysome analysis, single colonies were used to inoculate LB medium and grown overnight at 37°C with shaking. Overnight cultures were diluted 1:100 into fresh LB medium and grown for approximately 2 h at 37°C until exponential phase. Cultures were subsequently diluted to an OD_600_ of 0.02 and grown at the indicated temperatures (37°C, 25°C, or 20°C) until mid-log phase (OD_600_ 0.4–0.6). Translation was arrested by addition of chloramphenicol (100 μg mL⁻¹) for 5 min, followed by rapid cooling adding crushed ice. Cells were harvested by centrifugation at 4,000 ×g for 5 min at 4°C. Cell pellets were washed and resuspended in ice-cold wash buffer containing 10 mM Tris-HCl pH 7.5, 60 mM KCl, and 10 mM MgCl₂. Samples were normalised according to cell density and resuspended in lysis buffer containing wash buffer supplemented with 1 mg mL⁻¹ lysozyme, 1 mM PMSF, and RNase inhibitor. Cells were flash-frozen in liquid nitrogen and stored at −75°C until further processing. Cell lysates were generated by two freeze–thaw cycles consisting of freezing in liquid nitrogen and thawing at 10°C. Following the final thawing step, sodium deoxycholate was added to a final concentration of 0.6% together with RNase-free DNase I (0.05 U μL⁻¹). Lysates were incubated for 10 min at room temperature and clarified by centrifugation at 11,000 ×g for 15 min at 4°C. A total of 75 μL clarified lysate was loaded onto pre-cooled sucrose gradients and separated by ultracentrifugation using an SW41 rotor at 36,000 rpm for 2–6 hours at 4°C.

Gradients were analysed using a Piston Gradient Fractionator (Biocomp) while monitoring absorbance at 254 nm (A254). Resulting profiles were used to assess the relative distribution of free RNA, ribosomal subunits, 70S monosomes, and polysomes. For selected gradients, fractions were collected during profile acquisition to validate the identity of individual ribosomal peaks based on their rRNA content. Equal volumes of each fraction were mixed with an equal volume of 2× denaturing RNA loading dye and resolved on 2% (w/v) agarose gels in 0.5× TAE buffer stained with GelRed (Merck). Electrophoresis was continued until the 23S and 16S rRNA species were sufficiently separated, and gels were imaged using a Fusion Fx imaging system (Vilber Lourmat). The presence and relative distribution of 23S and 16S rRNA across fractions were used to support the assignment of 30S-, 50S-, and 70S-associated peaks.

Polysome profiles were aligned to the highest absorbance of the 70S monosome and plotted against fraction position using Prism (GraphPad), from the lowest to the highest sucrose concentration. Whole profiles and zoomed-in views of the ribosomal subunit and monosome regions were displayed to visualise ribosome biogenesis defects. Polysome-profiling experiments were independently repeated three times with similar results (N=3).

### Integrated rRNA synthesis, ribosome assembly and translation (iSAT) assay

Integrated rRNA synthesis, ribosome assembly and translation (iSAT) reactions were performed as described previously with minor adaptations^23,24^. Briefly, iSAT reactions were assembled at a final volume of 15 μL using the following final concentrations of components: 57 mM HEPES-KOH pH 7.6, 1.5 mM spermidine, 1 mM putrescine, 10 mM magnesium glutamate, 150 mM potassium glutamate, 1.6 mM ATP, 1.1 mM CTP, GTP, and UTP, 45.33 μg mL⁻¹ folinic acid, 227.5 μg mL⁻¹ tRNAs, 2 mM amino acid mixture, 42 mM phosphoenolpyruvate (PEP), 2 mM DTT, 0.33 mM NAD, 0.27 mM CoA, 4 mM oxalic acid, 1 μM T7 RNA polymerase, and 1 nM pJL1-sGFP reporter plasmid (kind gift from M. Jewett).

Premixed iSAT components were added to S150 cell extract to a final concentration of 33% (v/v). Ribosome assembly reactions contained 600 nM total ribosomal proteins (TP70), 4 nM pT7rrnB plasmid, and either 1.25 μM purified CsdA variants (WT, DQAD, ΔIDR1/2 or no CsdA spike-in). For translation-only control reactions, ribosome assembly components were omitted and replaced by 200 nM purified 70S ribosomes. Basal iSAT activity was assessed by performing reactions without addition of TP70.

Cell-free reactions were performed in technical triplicates in MicroAmp Fast Optical 96-well plates (Applied Biosystems) and incubated at 25 °C for 12 h in a StepOnePlus Real-Time PCR System (Applied Biosystems). sGFP fluorescence was measured every 10 min using the FAM/SYBR Green detection channel. For iSAT reactions performed at 37°C, data was collected every 5 min for 5 h. In addition to the iSAT reactions, GFP standards spanning 0.31–10 μM were measured under identical conditions to enable conversion of fluorescence signals into GFP concentrations.

For each GFP standard, fluorescence measurements acquired over time were averaged, and GFP concentrations were log_10_-transformed before fitting a linear regression of fluorescence intensity against log_10_-transformed GFP concentration. GFP concentrations in the iSAT reactions were determined at the indicated time point (4 or 6 h) by interpolating the measured fluorescence signal from this calibration curve and backtransforming the resulting concentration values. Interpolation was performed independently for each of the replicates. GFP concentrations are presented as the mean ± SD of the technical replicates for the controls and mean ± SD of two independent replicates for the main experiment (N=2). For each condition reaching a plateau the time to reach the half-maximum fluorescence of the curve was determined (T_50_) and mean ± SD of two independent replicates were plotted (N=2).

Significance was assessed using an ordinary one-way ANOVA with Fisher’s LSD test with a 95% confidential interval. Significance for selected pairwise comparisons are indicated as follows: ns, P ≥ 0.1234; P < 0.0332(*); P < 0.0021(**); P < 0.0002(***); P < 0.0001(****).

### Comparative genomics of the CsdA ortholog family Bacterial genome dataset construction

The GTDB was sampled^25^, and the bacterial species tree dated 02.07.2026 was used to construct a dataset of 290 bacterial genomes, selected to represent known bacterial taxonomic diversity at the “Phylum” level in a balanced manner. Genome quality was measured using the CheckM2 score^26^ reported on the NCBI databases^27^ to select the best genomes from each phylum, and were downloaded using their GCA accessions. 128 genomes had no gene annotations. To maintain consistency across all genomes, all 290 genomes were annotated with Pyrodigal 3.7.1^28^ to obtain CDS and protein sequences. This dataset was used to build a subset of the GTDB species tree for all downstream analyses.

### Bacterial genome annotation for optimal growth temperatures

GenomeSPOT, a set of machine-learning models that infer an organism’s growth conditions from genomic features, was used to infer optimal growth temperatures alongside other characteristics such as oxygen tolerance, pH, and salinity^29^. For each of the 290 genomes, the available assembly and the predicted proteomes were used under default conditions (genome_spot.genome_spot --models <models> --contigs <contigs.fna> --proteins <proteins.faa>). For each character, an optimum estimate is reported along with maximum/minimum tolerances, a per-prediction error estimate, and a novelty flag identifying genomes that are outliers relative to the training data. The predictions were compared with measured optimal growth temperatures taken from TEMPURA (25 genomes)^30^ and BacDive (13 genomes)^31^ for 38 genomes, matched using the NCBI taxonomy identifier.

### CsdA homolog detection and phylogeny

P0A9P6, the ATP-dependent DEAD-box helicase from E. coli, was used as the query to retrieve the top-scoring reciprocal-best-hit homologs from these 290 genomes. These homologs were aligned using MAFFT v7.525 under the “genafpair” strategy (mafft --anysymbol --maxiterate 1000 --bl 45 --genafpair --reorder --thread 2)^32^ and trimmed using trimAl v1.5.rev1 (build 2025-11-25)^33^ by discarding columns with over 60% gaps, and a gene tree was built with FastTree 2.2.0 (FastTree -spr 4 -mlacc 2 -slownni -gamma)^34^ to confirm orthology. This confirmed ortholog set was used to build a Hidden Markov Model (HMM) with hmmbuild under default parameters for the CsdA family with HMMER 3.4^35^, which was then used to search each of the 290 proteomes individually to detect diverged orthologs. For each genome, all hits with a best-1-domain E-value below 1E-10 were selected; when no hit satisfied this threshold, the top-scoring hit was retained, yielding 992 candidate homologs.

These homologs were aligned (mafft --anysymbol --auto --nofft --maxiterate 1000 --bl 30 --reorder --thread 2) and trimmed as above after removing 1 short sequence and 4 gappy sequences, giving an alignment of 987 sequences and 484 positions. A maximum-likelihood gene tree was inferred from this alignment using IQ-TREE v3.0.1^36^. To account for site-specific compositional heterogeneity, a FastTree tree of the same alignment was supplied as a guide tree to the infer the final tree by estimating posterior-mean site-frequency (PMSF) profiles approximating the LG+C20+F+R4 mixture model (iqtree -s <alignment> -m LG+C20+F+R4 - ft <guide_tree> -B 1000 -alrt 1000 -T 12)^37^, with branch support obtained from 1,000 ultrafast bootstrap replicates^38^ and 1,000 SH-aLRT replicates^39^.

The CsdA/CshA ortholog group was delineated from this maximum-likelihood tree. The clade encompassing the CsdA and CshA landmarks and their two closest paralog lineages was extracted and rooted on the CshB paralog clade; the immediately adjacent DbpA paralog clade was then excluded, delineating the CsdA/CshA ortholog group as 134 sequences across 121 genomes. A single representative per genome was retained for per-genome analyses.

### Sequence analyses

Intrinsically disordered regions (IDRs) were predicted for the sequences of the ortholog group and its two flanking paralog clades (DbpA and CshB) using metapredict v3 (3.0.2)^40^ with the default domain predictors. Non-standard amino acid residues were recoded to glycine, as required by metapredict. The C-terminal IDR, termed IDR2, was defined as the disordered domain with the largest end coordinate, and only when this coordinate lay within 30 residues of the C-terminus. As prediction was performed on the recoded sequences, the boundaries were transferred back to the original sequences; this was validated for the *E. coli* CsdA IDRs (IDR1: 439–487, IDR2: 554–629). The self-interaction parameter ε was calculated using the FINCHES^41^ implementation of the CALVADOS^42^ mean-field interaction model; ε was computed as a homotypic (self) interaction at a fixed reference temperature (i.e. temperature-independent), so that composition rather than the evaluation temperature determines ε. Where a genome contributed more than one tail-bearing sequence, the longest tail was used.

The sequence composition of the IDR2 tails was examined, focusing on the arginine-to-lysine balance R/(K+1), the density of the RG and GR dipeptides, and the fractional content of G, R, L, and aromatic residues. Within the ortholog group, the IDR2 tails were aligned using MAFFT (--auto --anysymbol), and per-column conservation was calculated as the product of column occupancy and the frequency of the most common non-gap residue. The 12 highest-scoring columns were taken as the motif anchor positions, and the consensus residue at each column was used to calculate the SLiM integrity^43^ of each ortholog IDR2 (the fraction of the 12 columns whose residues matched the consensus). Because this anchor-based score is defined on the ortholog-group consensus and is therefore not appropriate for cross-group comparison, an alignment-free RG-dipeptide density (the count of R–G adjacencies per residue) was computed in parallel as a non-circular cross-check.

To test whether the RGG/condensation tail is specific to the CsdA/CshA ortholog group, the tails were compared across the three clades delineated from the maximum-likelihood gene tree: the CsdA/CshA ortholog group, the intervening DbpA paralog clade, and the CshB paralog clade. Only the alignment-free metrics described above (RG-dipeptide density, R/(K+1), residue fractions, and ε) were used, deliberately avoiding the ortholog-consensus SLiM-integrity score to prevent circularity. Groups were compared using Mann–Whitney U tests and, for the subset of genomes carrying both a CshB and an ortholog-group tail, using paired Wilcoxon signed-rank tests, which control for organism, genomic GC content, and growth temperature.

### Ancestral-state reconstruction

Ancestral states were reconstructed on the maximum-likelihood ortholog gene tree, rooted on the CshB paralog clade. Because the low-complexity C-terminal tails cannot be reliably aligned across the group, an alignment-free approach was adopted in which ε and per-sequence character states were derived directly from each sequence. Presence or absence of the IDR2 tail was treated as a binary character and reconstructed by Fitch parsimony, from which the ancestral state and the numbers of independent gains and losses were obtained. The ancestral IDR2 ε was estimated as the Brownian-motion root state, computed as the phylogenetic generalised-least-squares mean under a Brownian covariance derived from the tree over the tail-bearing sequences. The number of independent origins of condensation propensity was estimated by binarising ε (attractive, ε < 0; repulsive, ε ≥ 0) and applying Fitch parsimony with ambiguous internal states resolved toward the repulsive state, counting the repulsive-to-attractive transitions along the tree. Reconstructions were performed in R with castor (maximum-parsimony ancestral states)^44^ and ape/phytools (Brownian-motion root state)^45,46^.

### Phylogenetic comparative analyses

Phylogenetic signal in IDR2 ε and in the tail arginine-to-lysine balance R/(K+1) was quantified by Blomberg’s K^47^, with significance assessed against 1,000 random tip permutations, and by Pagel’s λ^48^ estimated by maximum likelihood, using the covariance structure of the ortholog gene tree. Associations with optimal growth temperature were tested under explicit phylogenetic control on the GTDB species tree pruned to the genomes under analysis. Continuous relationships between optimal growth temperature and IDR2 tail length, ε, and — as a control for uniform genome streamlining — folded-core length (protein length minus the predicted IDRs) were evaluated by phylogenetic generalised least squares under a Brownian-motion covariance^49^. Presence or absence of the gene and of the IDR2 tail was modelled against optimal growth temperature by phylogenetic logistic regression (phyloglm, R package phylolm)^50,51^. Equivalent non-phylogenetic associations were assessed in parallel by Mann–Whitney U tests and Spearman rank correlations. Predicted optimal growth temperatures were used for the full genome set, while measured optimal growth temperatures (TEMPURA, treated as higher-confidence, and BacDive) were analysed both jointly and restricted to the TEMPURA subset; all tests were two-sided. Phylogenetic analyses were performed in R (ape, phytools, caper, phylolm, castor) (https://github.com/davidorme/caper).

## Data and material availability

All data associated with this study are presented in the main text or the supplementary material. *Raw imaging data and datasets associated with the phylogenetic analyses will be deposited in public repositories, with accession information provided upon publication.* All Materials used or generated in this study are commercially available or will be supplied upon reasonable request. All mass spectrometry proteomics data associated with this manuscript have been deposited to the ProteomicsXchange consortium via MassIVE (https://massive.ucsd.edu) with the accession number MSV000103029/PXD083276 (Dataset is currently private. Reviewers can access the dataset following the instructions on the website. Username: MSV000103029, Password: PCF).

## Code availability statement

Image Analysis code and phylogenetic analysis code is built using standard tools and available upon request. *All code will be deposited in public repositories upon editorial request or, at the latest, for publication*.

## Acknowledgments

We thank Marek Basler for providing access to the 3D-SIM microscope, and Melanie Engelin for their support in 3D-SIM imaging. We are grateful to Urs Jenal, Andreas Kaczmarczyk and the entire Jenal lab for very helpful discussions and continuously supporting us with their expertise in bacterial cell biology and genetics, sharing material and protocols. Thanks to the Biozentrum Imaging Core Facility in establishing and optimizing 3D-SIM imaging and the associated image-analysis workflow. We are grateful to the Biozentrum Proteomics Core Facility for support in setting up, performing, and analysing the proteomics experiments. We thank Timothy Sharpe and Tobias Mühlethaler from the Biozentrum Biophysics Facility for assistance with biophysical characterization assays and for valuable discussions. We thank Matej Siketanc for sharing the CellProfiler analysis pipeline used for the quantitative analysis of protein condensation, and Daan Overwijn for guidance in establishing protein purification procedures. We further thank Justin Meyer, Kareena Strunden, Tim Exnar, Mark Shepelev, Viktoria Hilgers and Jonas Bürki for their contributions to experimental and computational work during their student projects and apprenticeships. We are grateful to Philippe Van der Stappen and Ben Engel for their help and exciting project discussions. We thank the entire Hondele laboratory, especially Nicole Beuret, for stimulating discussions, ideas, and continuous support throughout this project. We also thank Peter Scheifele, Urs Jenal, Karsten Weis, and Miriam Linsenmeier for critically reading the manuscript and providing valuable feedback.

## Author contributions

M.J.G. and M.H. conceived and initiated the project. M.J.G. cloned and generated constructs and bacterial strains, purified recombinant proteins and performed the cellular experiments, including 3D-SIM imaging, growth assays and Western blots, as well as the biochemical and condensate experiments A.F. and M.J.G. jointly established and optimised the 3D-SIM imaging pipeline. S.A. contributed to cloning and strain generation, purified recombinant proteins, performed growth assays and carried out protein co-condensation assays. R.P. assisted with strain generation and supervised M.J.G. in establishing the respective methods.

K.D. performed and analysed the polysome-profiling experiments. N.S.Q. optimized and performed the iSAT assays. A.S. established and assisted with the mass-spectrometry experiments. K.R. and T.A.W. designed and conducted the comparative phylogenetic analyses. M.J.G., K.D., N.S.Q., K.R., A.S. and M.H. contributed to data analysis and interpretation. M.J.G. prepared the figures and data visualizations. M.J.G., T.A.W., O.D. and M.H. provided supervision, and T.A.W., O.D. and M.H. acquired funding supporting the work.

M.J.G. and M.H. wrote the original manuscript, with O.D. and T.A.W. contributing to its review and editing. All authors contributed to the interpretation of the data and conclusions of the study and reviewed the final manuscript. M.H. supervised and led the study.

## Competing interest declaration

The authors declare no competing or commercial interests.

**Supplementary information** Extended Data Figs. 1-8 Extended Data Tables 1-2

## Disclosure of AI use

For the comparative phylogenetic analyses, study design, ortholog delineation, and primary data generation were performed by K.R. Bacterial genome annotation, homolog searches, and maximum-likelihood analysis of the DEAD-box ATPase gene tree were also performed by K.R. Python/R scripts used for ancestral-state reconstruction, tail-loss/OGT analyses, and condensation analyses were developed with assistance from Claude Code using Claude Opus 4.8 (Anthropic), under the direction and review of K.R., who verified the code, analyses, and resulting outputs and takes full responsibility for their accuracy.

For image analysis, manual foci annotations for the training of foci detectors were performed by M.H. Code for the cell-segmentation pipeline, fine-tuning of Spotiflow foci detectors from manually annotated data, foci quantification, radial-enrichment analyses, and colocalisation-enrichment analyses was developed with assistance from Claude (Anthropic) via Claude Code, under the direction and review of M.H., who verified the analysis pipelines and resulting outputs and takes full responsibility for their accuracy.

Generative AI tools, including ChatGPT (OpenAI) and Claude (Anthropic), were also used during manuscript preparation to assist with language editing, sentence restructuring, and improving the clarity and concision of scientific writing. All AI-assisted text was critically reviewed and edited by the authors, who take full responsibility for the final manuscript. Consensus AI was additionally used to assist with literature searches and identification of potentially relevant publications. All cited literature was independently evaluated and verified against the original sources by the authors.

Study design, generation of underlying experimental data, data analysis unless otherwise indicated, determination of scientific conclusions, conceptualization and writing of the original draft were performed by the authors. All analyses, interpretations, citations, and final manuscript content were reviewed and approved by the authors.

## Corresponding authors

Correspondence to Maria Hondele.

## Extended Data Figures

**Extended Data Fig. 1.**
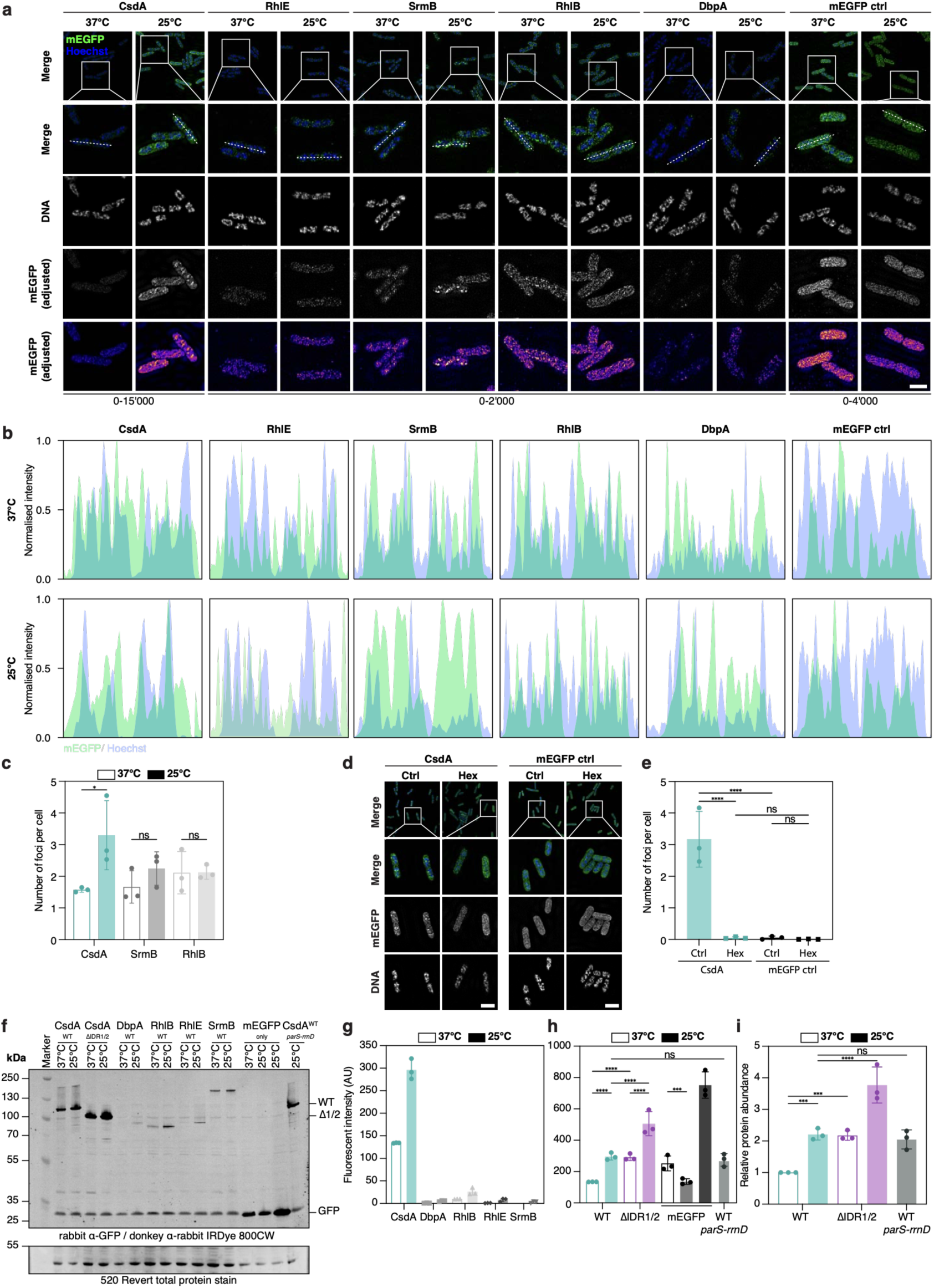
Cellular validation of cold-induced DEAD-box ATPase foci. **a,** 3D-SIM of the five endogenously mEGFP-tagged DDXs at 37 °C and 25 °C, displayed as maximum z projections for merged and individual channels. A soluble mEGFP-expressing WT strain was included as a negative control for focus formation. DNA was stained with Hoechst. Images were acquired using identical 3D-SIM settings (20% laser power and 40 ms exposure) and due to large differences in the DDX expression levels, brightness and contrast of the mEGFP channel was adjusted to show the optimal intensity range of each image as indicated. mEGFP is shown in black/white, as well as in fire highlighting foci (N=3). White dashed line identifies cells and pixels used to generate the intensity profiles in Extended Data Fig. 1b. For all 3D-SIM images scale bar: 2 µm. **b,** Normalised mEGFP and Hoechst fluorescence intensity profiles across representative cells shown in Extended Data Fig. 1a. **c,** Quantification of number of foci per cell for CsdA, SrmB and RhlB, in Fig. 1a and Extended Data Fig. 1a at 37 °C and 25 °C. CsdA was acquired using 5% laser power and 20 ms exposure; the other DDXs were acquired using 20% laser power and 40 ms exposure (N=3). Scare bar 2 µm. **d,** 3D-SIM of endogenously tagged CsdA-mEGFP at 25 °C before and after treatment with 5% 1,6-hexanediol for 5 min, displayed as maximum z projections for merge and individual channels. A soluble mEGFP-expressing WT strain was included as a negative control for focus formation. DNA was stained with Hoechst (N=3). Scale bar: 2 μm. **e,** Quantification of foci number per cell in untreated and 1,6-hexanediol-treated cells in Extended Data Fig. 2d. **f,** Representative anti-GFP immunoblot of the five endogenously mEGFP-tagged wild-type DDXs following growth at 37 °C and 25 °C, and the CsdA^ΔIDR1/2^ variant. Samples were normalised by OD_600_ before loading and 520 Rever total protein staining served as a loading control for quantification (N=3). **g,** Quantification of the five wild-type DDX signals from the immunoblot in Extended Data Fig. 2f. **h,** Quantification of mEGFP signal in the CsdA^WT^-mEGFP, CsdA^ΔIDR1/2^-mEGFP, soluble mEGFP and CsdA^WT^-mEGFP/*parS-rrnD* strain from the immunoblot in Extended Data Fig. 2f. **i,** Quantification of protein abundance in Extended Data Fig. 2f,h, relative to CsdA^WT^-mEGFP at 37 °C quantified by Western blot. **c,e,g,h,i,** Mean ± SD shown as bar graph, individual replicates shown as dots. Significance assessed by ordinary one-way ANOVA.

**Extended Data Fig. 2.**
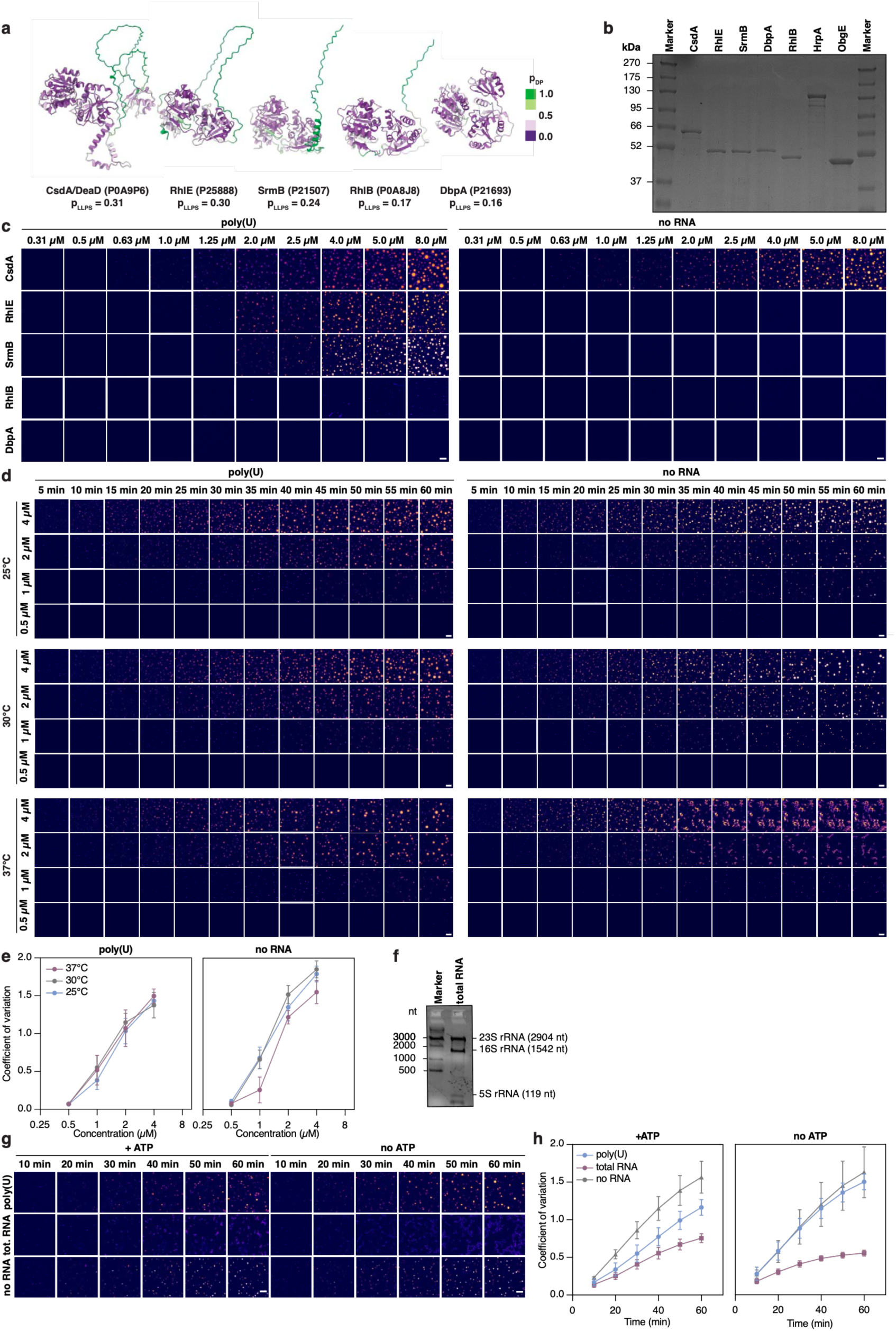
Biochemical validation of DEAD-box ATPase condensation. **a,** Residue-level predicted condensation propensity (pDB) calculated using FuzDrop superimposed on the AlphaFold-predicted structures of *E. coli* CsdA, SrmB, RhlE, RhlB and DbpA from the corresponding UniProt accession numbers are indicated. **b,** Coomassie stained SDS-PAGE analysis of 0.5 µg purified wild-type recombinant proteins used for the in vitro condensation experiments. Molecular-weight markers are indicated. **c,** Additional representative images of concentration-dependent condensation of recombinant CsdA, SrmB, RhlE, RhlB and DbpA in the absence or presence of poly(U) RNA imaged after 35 min at 25°C, corresponding to Fig. 1d,e. (N=3). Scale bar: 20 µm. **d,** Additional representative time- and concentration-resolved images of CsdA condensation at 25 °C, 30 °C and 37 °C corresponding to Fig. 1f-h (N=3). Scale bar: 20 µm. **e,** Concentration-dependent quantification by fluorescence coefficient of variation of CsdA at 25°C, 30°C and 37°C ± poly(U) (N=3). **f,** Agarose gel analysis of 1 µg total RNA isolated from *E. coli* MG1655 and used in the condensation assays. Positions of 23S, 16S and 5S rRNA are indicated relative to an RNA size marker. **g,** Additional representative time-resolved images of 2 µM CsdA incubated with poly(U), total *E. coli* RNA or no RNA in the presence or absence of ATP supplied with a creatine kinase-based ATP-regeneration system (CMK) at 25 °C, corresponding to Fig. 1i (N=3). Scale bar: 20 µm. **h,** Quantification of the ATP-dependent condensation kinetics of 2 µM CsdA condensation corresponding to Fig. 1i,j and Extended Data Fig. 2g in the presence of different RNAs. **e,h,** Mean ± SD.

**Extended Data Fig. 3.**
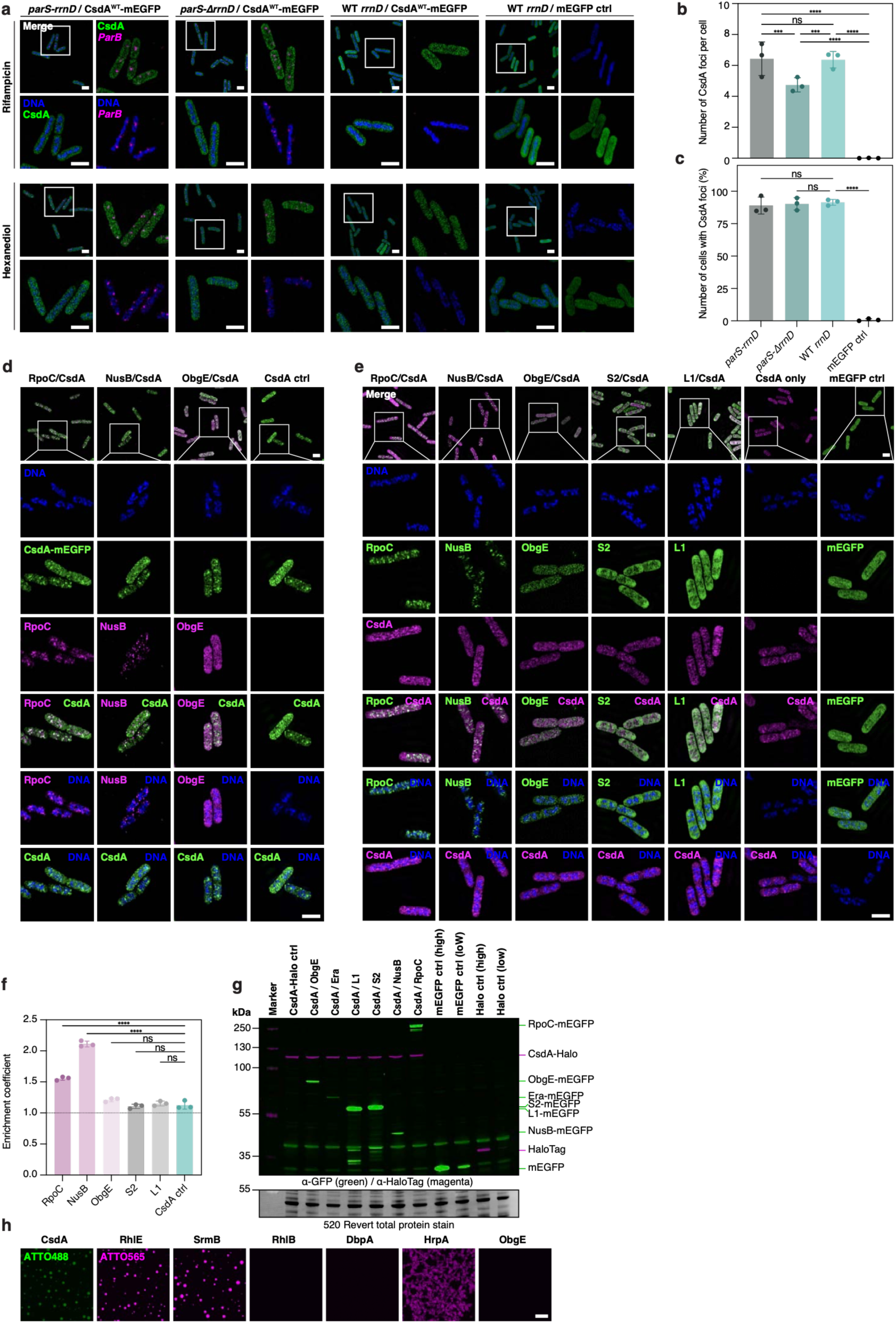
Additional *rrn* localisation and CsdA interaction analyses. **a,** Additional images of 3D-SIM corresponding to Fig. 2b-e, displayed as maximum z projections for merged and individual channels. CsdA^WT^-mEGFP and ParB/*parS*-labeled chromosomal loci with indicated perturbation using rifampicin or 1,6-hexanediol following growth at 25°C. DNA was stained with Hoechst (N=3). Scale bar: 2 µm. **b,c,** Number of CsdA foci per cell, and percentage of cells containing CsdA foci of cells containing CsdA foci in the *parS-rrnD*, *parS-ΔrrnD* and control strains corresponding to Fig. 2b. **d,** Additional separate fluorescence channels corresponding to Fig. 2f showing DNA stained with Hoechst, CsdA^WT^-mEGFP together with plasmid-expressed mKOk-tagged RpoC, NusB or ObgE and different merge combinations following growth at 25°C (N=3). Scale bar: 2 μm. **e,** 3D-SIM of endogenously tagged CsdA^WT^-Halo together with the indicated endogenously mEGFP-tagged ribosome biogenesis associated factors following growth at 25 °C. CsdA-Halo was labelled with Janelia Fluor 549 HaloTag. DNA was stained with Hoechst. (N=3). Scale bar: 2 μm. **f,** Enrichment of endogenously tagged ribosome biogenesis associated factors within CsdA-Halo condensates in Extended Data Fig. 3e. **g,** Anti-GFP and anti-HaloTag fluorescent immunoblot analysis of the CsdA^WT^-Halo parental strain and CsdA^WT^-Halo, POI-mEGFP double-tagged strains used in Extended Data Fig. 3e,f. Samples were normalised by OD_600_ and 520 Revert total protein staining served as loading control. Molecular-weight markers are indicated (N=3). **h,** Representative in vitro condensation images of 2 µM of the indicated protein alone, serving as controls for the in vitro co-condensation experiments shown in Fig. 2h,j (N=3). Scale bar: 20 µm. **b,c,f** Mean ± SD shown as bar graph, individual replicates shown as dots. Significance assessed by ordinary one-way ANOVA.

**Extended Data Fig. 4.**
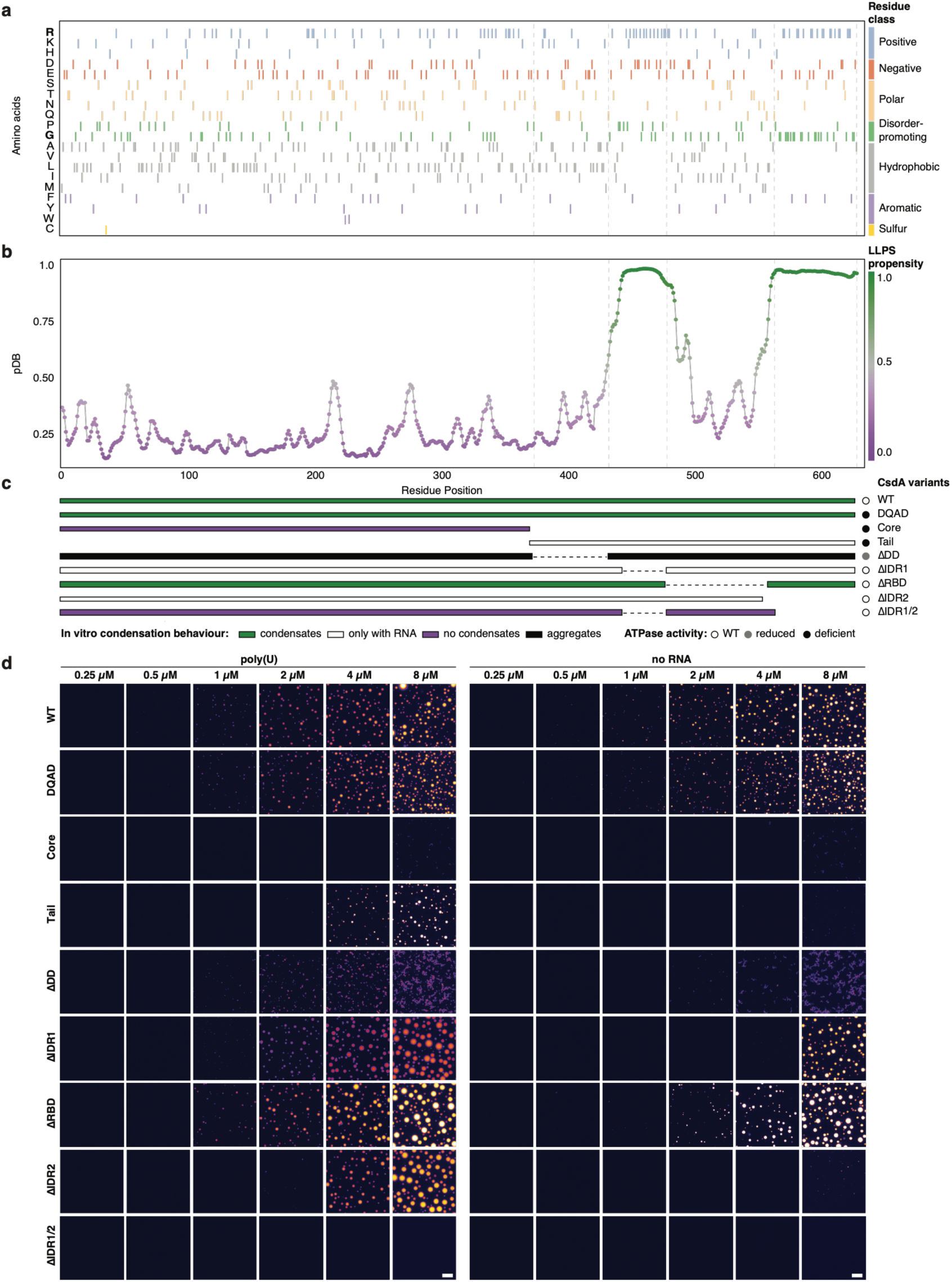
Biochemical dissection of CsdA condensation determinants. **a,** Amino-acid composition across the CsdA sequence aligned to the CsdA domain architecture indicated by horizontal grey dashed-lines. Residues are grouped according to physicochemical properties; positive, negative, polar, disorder-promoting, hydrophobic, aromatic and cysteine containing. **b,** Residue-level condensation propensity (pDB) predicted by FuzDrop along the CsdA sequence from the UniProt number P0A9P6. **c**, CsdA variants used to dissect condensation and enzymatic activity and experimentally determined phenotypes indicated. **d,** Additional representative images of wild-type CsdA and mutant variants across concentrations of 0.25–8.0 μM in the ± poly(U), corresponding to Fig. 3b,c (N=3) Scale bar: 20 μm.

**Extended Data Fig. 5.**
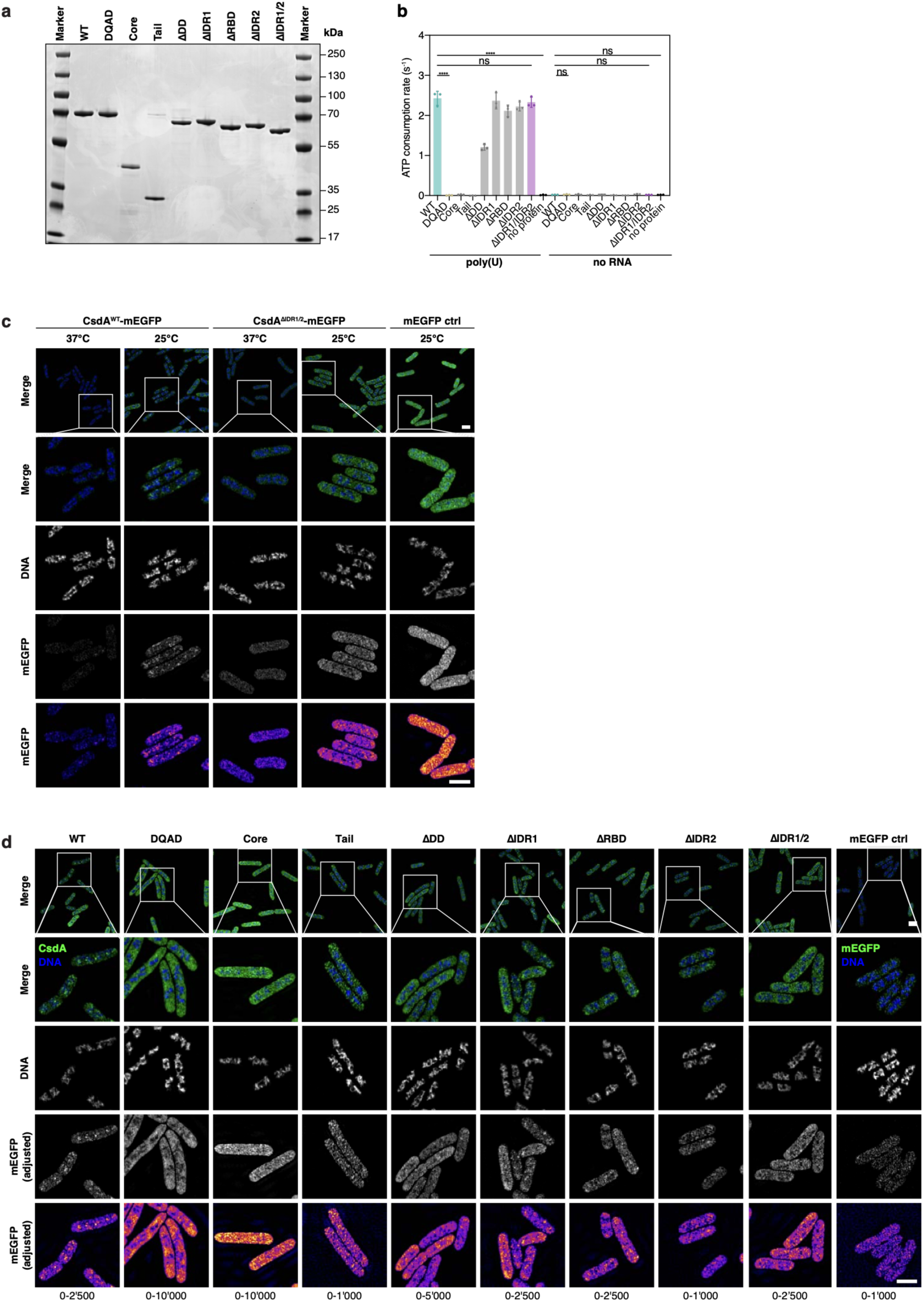
Cellular phenotypes of CsdA condensation mutants. **a,** Coomassie stained SDS-PAGE analysis of 0.5 µg purified wild-type CsdA and mutant variants used in Fige. 3 and 4. Molecular-weight markers are indicated. **b,** ATPase activity of wild-type CsdA and indicated variants in the ± poly(U), complementing the RNA-stimulated measurements in Fig. 3d (N=3). Mean ± SD shown as bar graph, individual replicates shown as dots. Significance assessed by ordinary one-way ANOVA **c,** Separate channels of 3D-SIM displayed as maximum z projections corresponding to Fig. 3e of CsdA^WT^-mEGFP and CsdA^ΔIDR1/2^-mEGFP following growth at 37°C or 25°C. DNA was stained with Hoechst. Images were acquired using identical 3D-SIM settings and displayed using matched brightness and contrast (N=3). Scale bar: 2 µm. **d,** Separate channels of 3D-SIM displayed as maximum z projections corresponding to Fig. 3f for endogenously tagged wild-type CsdA and the indicated variants following growth at 25°C. DNA was stained with Hoechst. Images were acquired using identical 3D-SIM settings (5% laser power and 20 ms exposure) and due to large differences in variant expression levels, brightness and contrast of the mEGFP channel was adjusted to show the optimal intensity range of each image as indicated. mEGFP is shown in black/white, as well as in fire highlighting foci (N=3). Scale bar: 2 µm.

**Extended Data Fig. 6.**
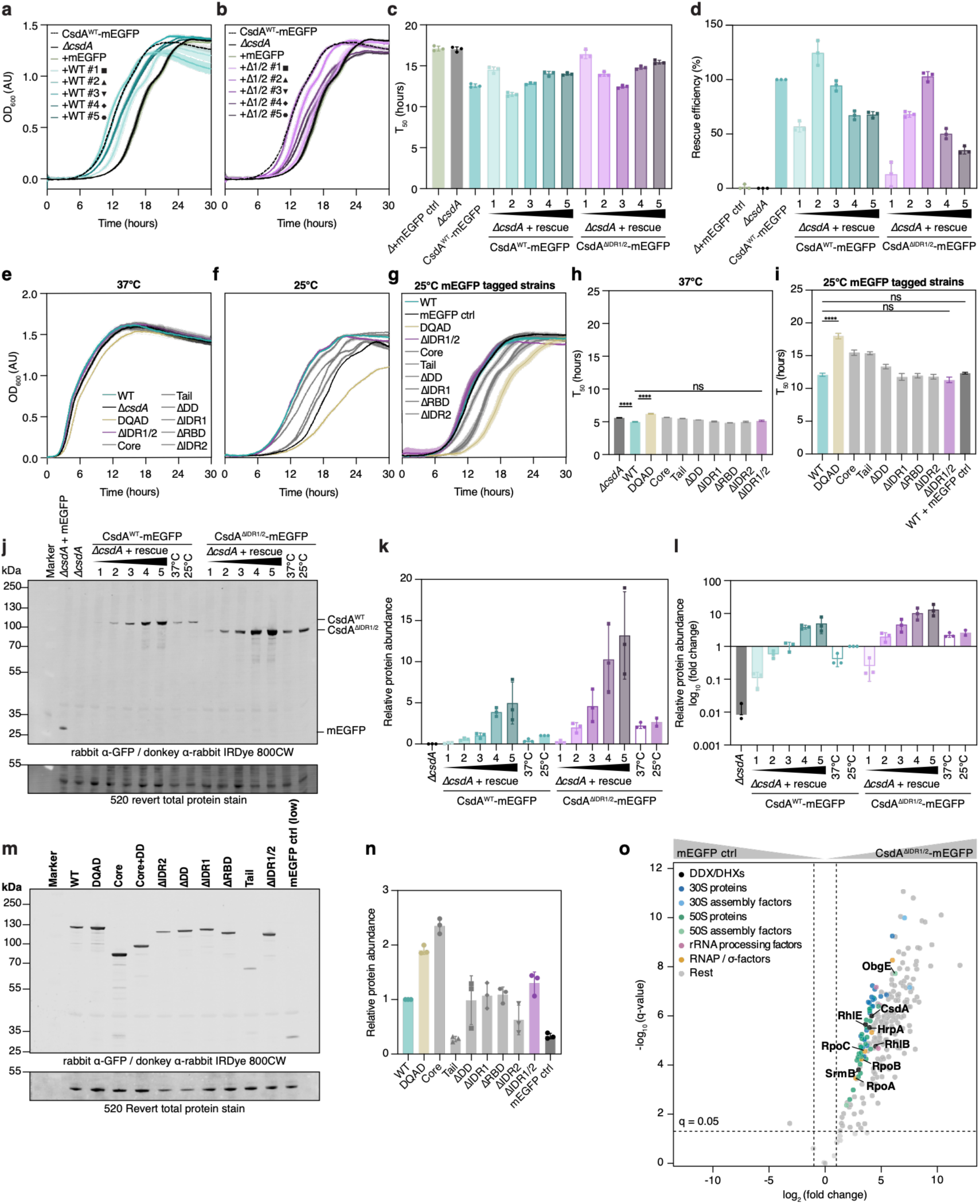
Expression-dependent rescue and CsdA interactome analyses. **a,b,** Rescue growth curves of Δ*csdA* cells expressing wild-type CsdA (a) or CsdA^ΔIDR1/2^ (b) from promoters of graded strength at 25 °C (increasing from 1 to 5, WT in shades of teal, ΔIDR1/2 in shades of purple). CsdA^WT^-mEGFP (black, dashed), Δ*csdA* (black, solid) and Δ*csdA* expressing soluble mEGFP (green, solid) from plasmid were included as positive (WT) and negative (no rescue) controls, respectively (N=3). **c,** Quantification of time to half-maximal OD_600_ (T_50_) derived from the rescue growth curves at 25°C in Extended Data Fig. 6a,b. **d,** Quantification of relative rescue efficiency in percent derived from the T_50_ of the rescue growth curves at 25 °C in Extended Data Fig. 6c. Based on 100% rescue of CsdA^WT^-mEGFP control and no rescue of the Δ*csdA* negative control. **e,** Growth curves of strains expressing endogenous wild-type CsdA and the indicated CsdA variants at 37°C (N=3). **f,** Growth curves of strains expressing endogenous wild-type CsdA and the indicated CsdA variants at 25°C (N=3). **g,** Growth curves of strains expressing endogenous mEGFP-tagged wild-type CsdA and the indicated CsdA variants at 25°C (N=3). **h,** Quantification of T_50_ for the growth curves of endogenous wild-type CsdA and the indicated CsdA variants at 37 °C in Extended Data Fig. 6e. **i,** Quantification of T_50_ for the growth curves of endogenously mEGFP tagged wild-type CsdA and the indicated CsdA variants at 25 °C in Extended Data Fig. 6g (N=3). **j,** Representative anti-GFP immunoblot used to calibrate cellular abundance of wild-type CsdA and CsdA^ΔIDR1/2^ expressed in the Δ*csdA* strain in the rescue experiment Extended Data Fig. 6a-d at 25 °C. Samples were normalised by OD_600_ before loading and total protein staining served as a loading control for quantification (N=3). **k,l,** Quantification of protein abundance in Extended Data Fig. 6j expressed relative to wild-type CsdA at 25 °C. (k) linear and (l) log_10_-transformed complementing Fig. 3j. **m,** Representative anti-GFP immunoblot of endogenously mEGFP-tagged wild-type CsdA and mutant variants following growth at 25 °C in Extended Data Fig. 6g,i (N=3). **n,** Quantification of protein abundance in Extended Data Fig. 6m, expressed relative to wild-type CsdA at 25°C. **o,** AP-MS of CsdA^ΔIDR1/2^-mEGFP interaction partners following cold adaptation compared to the mEGFP control. DDXs, ribosomal proteins and ribosome biogenesis factors, RNA processing factors, RNA polymerase subunits (RNAP) and sigma factors highlighted. Log_2_(FC) and q-values represent the mean of three independent biological replicates (N=3). Dashed lines: q=0.05 and |log_2_(FC)|=1 threshold. **a-i,k,l,n,** Mean ± SD. Significance accessed by ordinary one-way ANOVA.

**Extended Data Fig. 7.**
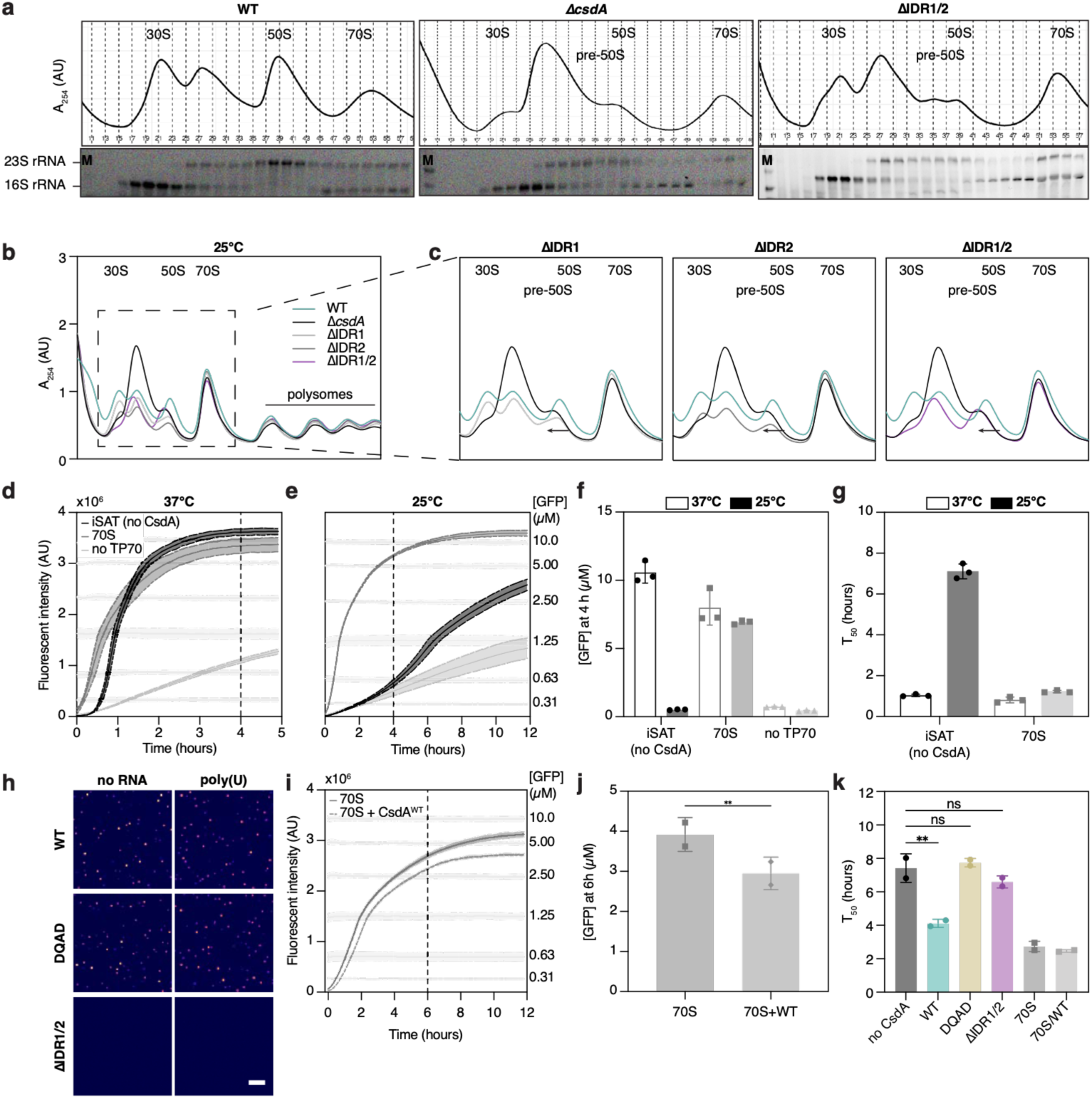
Additional ribosome assembly and iSAT analyses. **a,** Representative sucrose-gradient profiles from cells (WT, Δ*csdA*, or ΔIDR1/2) grown at 25 °C together with agarose gel analysis of RNA isolated from corresponding gradient fractions to identify pre-30S/30S, pre-50S/50S and 70S fractions. Positions of the 30S, 50S and 70S peaks are indicated based on the agarose gel, where positions of 16S and 23S rRNAs are highlighted (N=3). **b,** Representative sucrose-gradient profiles of cells expressing wild-type CsdA, CsdA^ΔIDR1^, CsdA^ΔIDR2^, CsdA^ΔIDR1/2^ from the endogenous locus or lacking CsdA (Δ*csdA*) following growth at 25 °C (N=3). **c,** Enlarged views of the profiles in Extended Data Fig. 7b, highlighting altered ribosome assembly intermediates and rescue potential of the individual IDR deletions. **d,e,** Representative GFP reporter accumulation in iSAT control reactions performed without exogenous CsdA, or addition of mature 70S ribosomes or without TP70 at 37 °C d, or 25 °C e, in technical triplicates. The dashed-line indicates the time-point used for GFP quantification in Extended Data Fig. 7f. **f,** Quantified GFP concentration after 4 h in iSAT control reactions performed at 37 °C and 25 °C (Extended Data Fig. 7d,e). **g,** Quantified time to half-maximal GFP fluorescence (T_50_) for the control reactions at 37 °C and 25 °C (Extended Data Fig. 7d,e). **h,** Representative images of in vitro condensation of 1.25 µM CsdA^WT^, CsdA^DQAD^ and CsdA^ΔIDR1/2^ in iSAT mimicking buffer conditions ± poly(U) at 25 °C (N=3). Scale bar: 20 μm. **i,** Representative GFP reporter accumulation in iSAT control reactions performed with mature 70S ribosomes ± CsdA^WT^ at 25 °C in technical triplicates. Additional data from Fig. 4e. The dashed-line indicates the time-point used for GFP quantification in Extended Data Fig. 7j (N=2). **j,** Quantified GFP concentration after 6 h in iSAT control reactions performed at 25°C in Extended Data Fig. 7i. **k,** Quantified time to half-maximal GFP fluorescence (T_50_) iSAT reactions performed at 25 °C without added CsdA (no CsdA) or supplemented with wild-type CsdA, DQAD or ΔIDR1/2, corresponding to Fig. 4e (N=2). **d-g,i-k,** Mean ± SD. **f-g,** individual dots represent technical replicates, **j-k,** individual dots indicate biological replicates. Significance assesses by ordinary one-way ANOVA.

**Extended Data Fig. 8.**
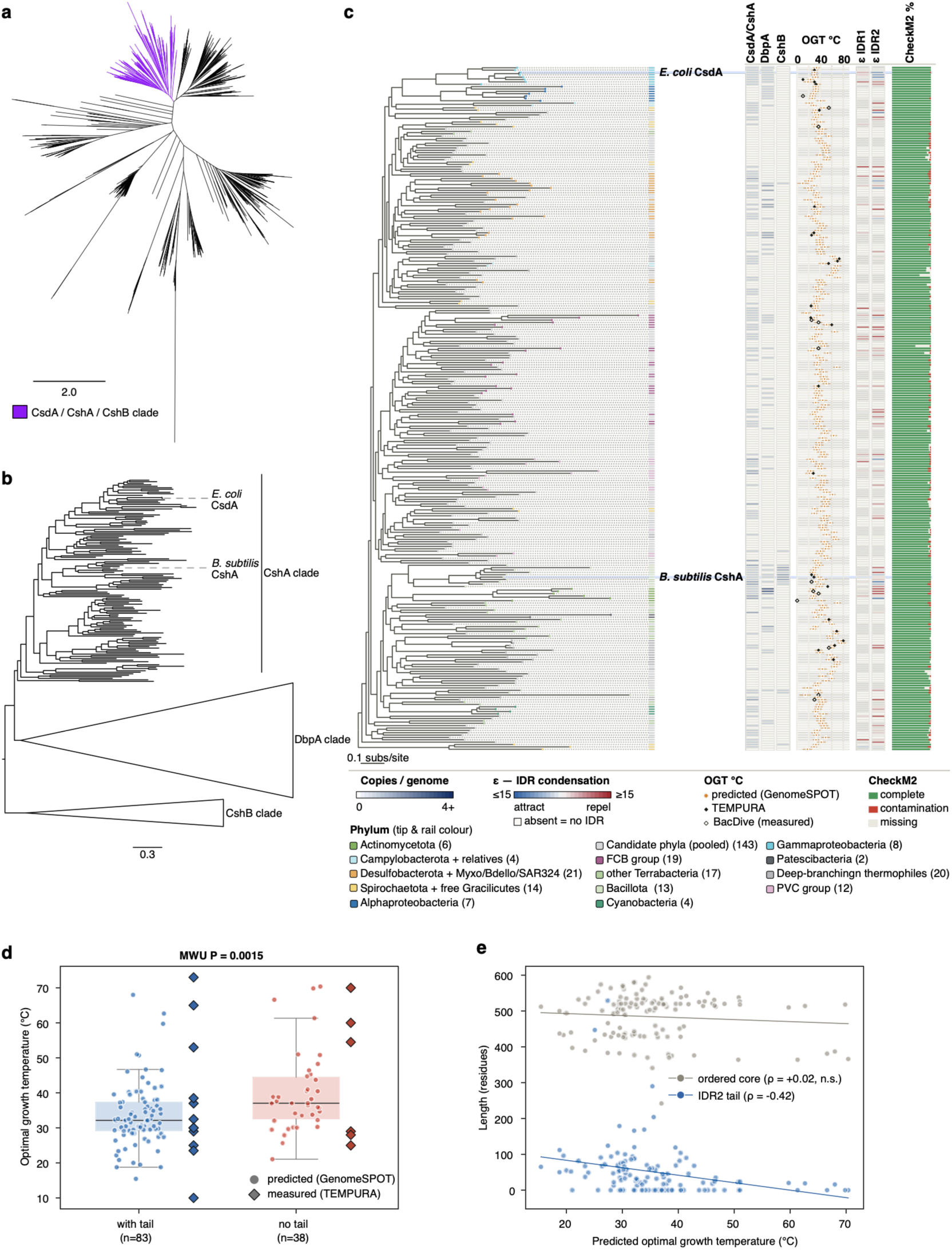
Evolutionary conservation of the CsdA/CshA condensation module. **a,** Full unrooted gene tree (LG+C20+F+R4, 1000 ultrafast bootstraps, 1000 SH-aLRT replicates, bootstraps not shown for convenience) of DEAD-box helicases as extracted using the CsdA HMM, with the clade containing CsdA/CshA/CshB shaded. Tree scale reported in number of substitutions per site. **b,** Phylogenetic profile of different clades of the CsdA family gene tree, predicted OGT, IDR1 and IDR2 epsilon values, CheckM2scores for the 290 bacterial genomes. Species tree tips are coloured based on the lineage/group to which the species belongs. The phylogenetic profile is coloured as a gradient indicating number of copies (darker blue indicates more copies). The IDR ε profile is coloured based on condensation propensity (blue indicates attractive i.e. condensation-prone, red indicates the opposite). Diamonds in the OGT graph indicate experimentally measured OGTs (filled diamonds are TEMPURA measurements, empty diamonds are BacDive measurements). Tree scale reported in number of substitutions per site. **c,** Structure of the CsdA unrooted subtree. Clades were annotated based on the presence of either the *E. coli* protein or the *Bacillus subtilis* (*B. subtilis*) protein. Tree scale reported in number of substitutions per site. **d,** Orthologs lacking an IDR2 tail come from significantly warmer organisms (predicted optimal growth temperature; Mann–Whitney U P = 0.0015; medians 37.0 vs 32.1°C). Diamonds show directly measured optima (TEMPURA), which trend the same way. **e,** IDR2 tail length decreases with optimal growth temperature (Spearman ρ = −0.42) whereas the ordered core (protein length minus predicted IDRs) is invariant (ρ = +0.02, ns), indicating tail-specific loss rather than genome-wide streamlining. The significant axis uses composition-derived predicted temperatures; the effect also survives phylogenetic control (PGLS: presence P = 0.019, length P = 0.005).

